# The spatio-temporal program of liver zonal regeneration

**DOI:** 10.1101/2021.08.11.455924

**Authors:** Shani Ben-Moshe, Tamar Veg, Rita Manco, Stav Dan, Aleksandra A. Kolodziejczyk, Keren Bahar Halpern, Eran Elinav, Shalev Itzkovitz

## Abstract

The liver carries a remarkable ability to regenerate rapidly after acute zonal damage. Single-cell approaches are necessary to study this process, given the spatial heterogeneity of multiple liver cell types. Here, we use spatially-resolved single cell RNA sequencing (scRNAseq) to study the dynamics of mouse liver regeneration after acute acetaminophen (APAP) intoxication. We find that hepatocytes proliferate throughout the liver lobule, creating the mitotic pressure required to repopulate the necrotic pericentral zone rapidly. A subset of hepatocytes located at the regenerating front transiently up-regulate fetal-specific genes, including Afp and Cdh17, as they reprogram to a pericentral state. Zonated endothelial, hepatic-stellate cell (HSC) and macrophage populations are differentially involved in immune recruitment, proliferation and matrix remodeling. We observe massive transient infiltration of myeloid cells, yet stability of lymphoid cell abundance, in accordance with global decline in antigen presentation. Our study provides a resource for understanding the coordinated programs of zonal liver regeneration.

## Introduction

The liver is a highly heterogeneous organ. Hepatocytes and supporting non-parenchymal cells operate in repeating hexagonally-shaped anatomical units termed ‘liver lobules’. Blood enters the lobules at their corners, termed ‘portal nodes’ and flows inwards through sinusoidal channels into draining central veins. This polarized blood flow, in conjunction with sequential consumption and secretion of hepatocytes generates gradients of oxygen, nutrients and hormones. The gradients of blood-borne factors and additional morphogen gradients create a highly heterogeneous microenvironment. As a result, cells at different lobule coordinates exhibit notable differences in gene expression, a phenomenon termed ‘liver zonation’ (Ben-Moshe & Itzkovitz, 2019; Gebhardt, 1992; Jungermann & Keitzmann, 1996). About half of the hepatocyte genes are zonated, with processes such as gluconeogenesis and protein secretion allocated to the periportal hepatocytes and other processes such as drug detoxification allocated to pericentral hepatocytes (Halpern et al., 2017). Liver endothelial cells (Halpern et al., 2018; Inverso et al., 2021) and hepatic stellate cells (Dobie et al., 2019) exhibit similar broad functional zonation patterns along the lobule radial axis. It is unclear how this spatial heterogeneity affects liver pathology and regeneration processes.

The liver exhibits robust regenerative capacity (Michalopoulos & DeFrances, 1997). In response to acute doses of drugs, such as acetaminophen (APAP, Bhushan & Apte, 2019; Mossanen & Tacke, 2015) or CCl_4_ (Recknagel, 1967), pericentral hepatocytes that attempt to detoxify these foreign substances are overwhelmed with toxic intermediates and die. The remaining liver tissue enters a regenerative mode, leading to rapid healing and replacement of the damaged lobule layers (Yanger et al., 2014). Zonal regeneration involves a coordinated set of processes that have to be carried out in the right place and time. Dead hepatocytes need to be efficiently cleared, while preventing induction of the adaptive immune system exposed to the multiple new available antigens. Extra-cellular matrix needs to be constructed rapidly to support the tissue scaffold, yet disintegrated once new cells are formed, to prevent lasting fibrosis (Adler et al., 2020; Kisseleva & Brenner, 2006). Most importantly, hepatocytes originating in lobule zones with profoundly different expression signatures need to be rapidly generated and reprogrammed to take over the critical pericentral hepatocyte functions. Single cell approaches have been used to study non-parenchymal cells during the acute phase of APAP intoxication at 20hrs after damage (Kolodziejczyk et al., 2020), yet the spatial and temporal dynamics of the coordinated response of all liver cell types across the entire zonal regeneration process have not been explored.

## Results

### A single cell atlas of zonal regeneration after APAP

To study zonal regeneration, we injected mice with an acute dose of 300mg/kg APAP (Figure 1A) and sampled livers at different time points after injection. We observed massive pericentral necrosis peaking at 48h after APAP (Figure 1B), apparent from the lack of hepatocyte nuclei. The damaged pericentral areas contracted at 72h with the appearance of new pericentral hepatocytes, which completely replaced the necrotic tissue after 96h (Figure 1B). To identify the molecular details of the regeneration process we performed bulk RNAseq and scRNAseq at multiple time points (Figure 1A, Figure S1). Our bulk RNAseq included 21 mice sacrificed at seven time points after APAP injections, starting at 6h and up to 1month following injury (three mice per time point), and eight controls injected with saline at four different time points (Table S1). Our scRNAseq cell atlas included 20,587 cells collected from 19 mice at 6 time points – control, 24h, 48h, 72h, 96h and 1w following injury. The single cells clustered into 11 cell types, each exhibiting distinct marker genes (Figure 1C-D, Figure S1A, Table S2). These cell types included hepatocytes, endothelial cells, HSCs, Kupffer cells, cholangiocytes, monocytes, macrophages, plasmocytoid dendritic cells (pDCs), conventional dendritic cells (cDCs), B cells and a cluster of T cells and NK cells.

**Figure 1.**
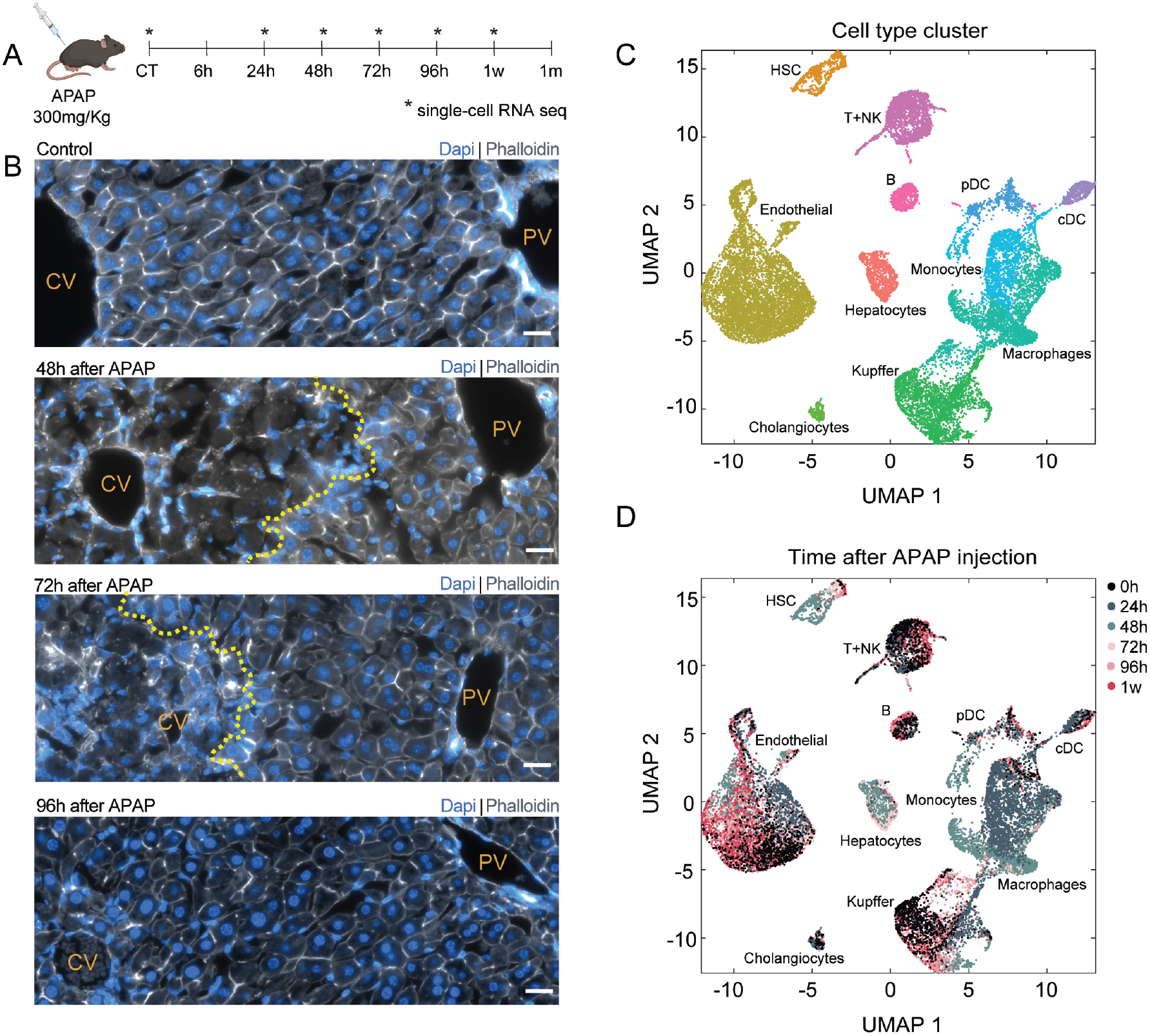
Acute dose of APAP induces liver damage and regeneration. (A) A schematic of the experimental design. Mice were injected with 300mg APAP / 1kg body weight. Livers were harvested for bulk sequencing at 6h, 24h, 48h, 72h, 96h, 1 week and 1 month after injection (3 mice per time point), as well as saline injected controls at different time points (2 mice per time point). Livers were dissociated for single cell sequencing at 24h, 48h, 72h, 96h, 1 week and non-treated controls (2-4 mice for each time point, marked with asterisks). Mouse injection illustration was created with BioRender.com. (B) Images of liver lobules at different time points following APAP injection. CV – central vein, PV – portal vein. Yellow dashed line mark the borders of the damaged areas. Cell nuclei are stained with Dapi (blue). Cell membranes are stained with Phalloidin (grey). Scale bar – 20*μm*. (C) Uniform Manifold Approximation and Projection (UMAP) visualization of the integrated data of all 20,587 cells from 6 time points (n=19 mice). Cells are colored by their cell type annotation. (D) UMAP visualization of the integrated data. Cells colored by the time following APAP injection.

Principal component analysis (PCA) of the bulk RNAseq showed divergence in liver gene expression at 6h, 24h and 48h, with a reversion to the control signature by 96h (Figure S1B, Table S1). We used computational deconvolution of the bulk data using our scRNAseq cell-type signatures (Newman et al., 2019) to estimate the relative abundance of each cell type at different time point (Methods). Hepatocyte abundance significantly declined at 48h after APAP injection, in line with the substantial necrosis of pericentral hepatocytes (Figure 1B), and reverted back to the control proportions by 96h (Figure S1C). In contrast, HSCs, monocytes and macrophages increased in abundance, peaking at 24h-48h after APAP injection before declining back to control levels (Figure S1C). In accordance with this peak expansion at 48h, our scRNAseq atlas revealed a peak in the proportion of proliferating cells for monocytes, macrophages and HSCs at 48h (Figure S1D). Notably, Kupffer cells and endothelial cells also exhibited a peak in proliferation at 48h, yet did not show a corresponding increase in abundances (Figure S1C, S1D, Table S3). This could indicate that, in addition to hepatocytes, some Kupffer and endothelial cells may have died during the acute injury phase and have been replenished by proliferation of the remaining cells, as previously demonstrated for Kupffer cells following APAP (Zigmond et al., 2014). Our analysis therefore expose substantial changes in liver gene expression and in proportions of multiple liver cell types, yet a notable rapid reversion to control levels already 4 days after damage. We next set out to characterize the dynamic molecular programs of this regeneration process for distinct liver cell types.

### Hepatocytes exhibit zonal reprogramming and broad proliferation across the liver lobule

Pericentral and periportal hepatocytes exhibit distinctly different gene expression in unperturbed livers (Halpern et al., 2017). As observed for the bulk RNAseq of the complete liver transcriptome (Figure S1B), the transcriptomes of hepatocyte-specific genes diverged from controls at early time points but reverted to control levels by 96h post APAP injection (Methods, Figure 2A). Our hepatocyte single cells data consisted of 2,770 cells from control (Methods), 48h and 72h following APAP injection (Figure 2B). We used the single-cell expression patterns of a large set of hepatocyte zonated landmark genes (Droin et al., 2021) to infer the zonation coordinates along the liver lobule axis for individual sequenced hepatocytes (Methods, Figure 2C-F). Subsequently, we binned cells from each time point to three zones: pericentral, mid-lobule and periportal (Table S4). As expected, hepatocytes with a pericentral signature were depleted at 48h. The distribution of zonation coordinates approached the pre-injury pattern at 72h (Figure 2F). We used single molecule fluorescence in-situ hybridization (smFISH) to demonstrate that classic zonated genes such as the pericentral Glul and Cyp2e1 and the periportal Ass1 assumed their pre-injury zonation patterns at 96h after damage (Figure 2G-H). Notably, newly formed pericentral hepatocytes exhibited significantly higher ploidy levels, an increase that was more prominent than in periportal hepatocytes (Figure 2I-J). Our analysis therefore show that hepatocyte re-assume their zonal molecular identity 4 days after acute liver damage.

**Figure 2.**
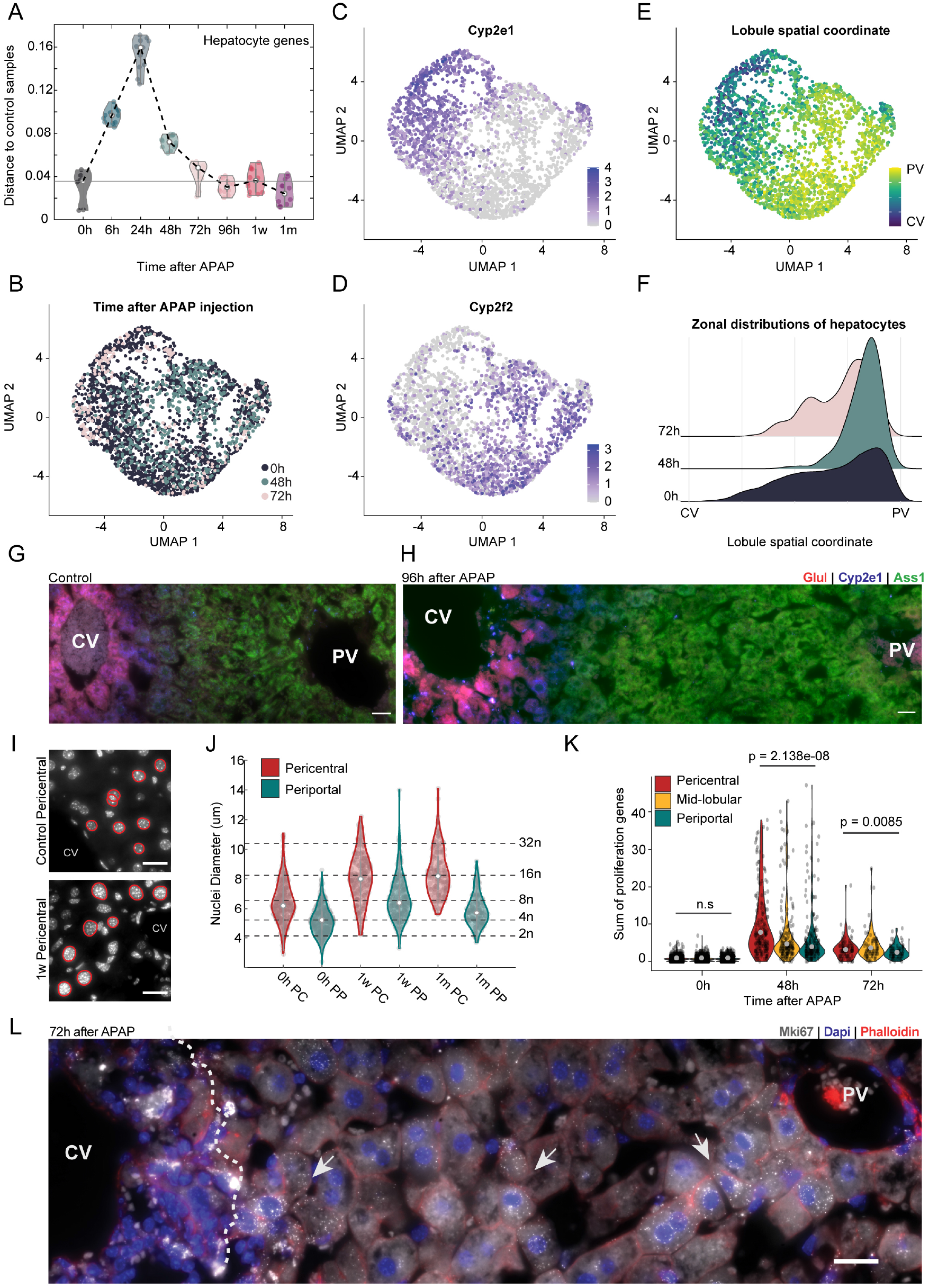
Hepatocytes from all zones proliferate and re-establish zonation. (A) Spearman correlation distances between hepatocyte-specific genes for pairs of control samples (n = 4) and mice at different time points (n = 3 mice per time point) after APAP injection. Horizontal line denotes median distance of control-control pairs. White dots are median distances for each time point. (B) UMAP visualization of hepatocytes (n = 2,770 cells), colored by time after APAP injection. (C)UMAP visualization of hepatocytes colored by the expression of the centrally zonated gene Cyp2e1. (D) UMAP visualization of hepatocytes colored by the expression of the portally gene Cyp2f2. (E) UMAP of hepatocytes colored by their inferred lobule spatial coordinate, ranging from CV – cells closest to central vein, to PV – cells closest to portal veins. (F) Distributions of lobule spatial coordinates of hepatocytes at each time point. (G-H) smFISH of a liver lobule showing 3 zonated genes: pericentral Glul (red) and Cyp2e1 (blue) and periportal Ass1 (green). CV – central vein. PV – portal vein. Scale bar - 20*μm*. Shown are examples of a control lobule (G) and a lobule 96h after APAP (H). (I) Dapi images of pericentral hepatocyte nuclei in control (top) and 1 week (bottom) following APAP injection. CV – central vein. Scale bar - 20*μm*. Represented circumferences are circled in red. (J) Quantification of nuclei diameters from H&E scans of hepatocytes located pericentral (red) and periportal (green) hepatocytes, in livers from control, 1w and 1m post APAP injection. Grey dots represent single nuclei, white dots are median diameters. Dashed horizontal lines correspond to mean diameters of different ploidy classes (Tanami et al., 2017). n = 1,518 nuclei, pooled from 2 mice per time point. (K) Sum of expression of proliferation genes (Methods) in hepatocytes from the different time points, grouped by their zone – pericentral (red), mid-lobular (yellow) or periportal (green). Black dots represent single cells, grey dots denote group median. Significance levels calculated using Kruskal-Wallis tests. (L) smFISH of a liver lobule 72h following APAP administration showing Mki67 single transcripts (grey dots). Dashed white line marks the damage border, arrows point to representative Mki67+ proliferating cells. Cell nuclei are stained with DAPI (blue) and membranes are stained with phalloidin (red). Scale bar - 20*μm*.

To identify the source of the regenerating cells, we analyzed the zonal distributions of proliferating hepatocytes. Hepatocytes proliferated throughout the lobule axis, with a bias towards the pericentral damage zone at 48h and towards the mid-lobule zones at 72h (Figure 2K). We used smFISH to validate the existence of Mki67+ hepatocytes at the mid-lobular and portal zones, at large distances from the damaged tissue area (Figure 2L). The broad proliferation of hepatocytes throughout the lobule axis may serve to generate an increased mitotic pressure along the hepatic plates that may contribute to rapidly replacing the cells at the damaged area. Importantly, this mitotic pressure brings mid-lobular hepatocytes into the pericentral zone, requiring reprogramming their transcriptional states (Figure 2F, H).

### Interface hepatocytes up-regulate fetal programs and exhibit a mesenchymal shape

To explore the cellular processes involved in the reprogramming of the mid-lobular hepatocytes into pericentral cell states, we used our scRNAseq measurements to compare the transcriptomes of the most pericentral hepatocytes at 48h and 72h and control hepatocytes at matching corresponding zones (Methods). These hepatocytes constituted the cells at the interface between the damaged and non-damaged zones, which we expected would undergo profound changes in zonal gene expression. We therefore termed these cells ‘interface hepatocytes’. We found that interface hepatocytes exhibited a distinct expression signature, consisting of genes that are expressed in fetal livers and in hepatocellular carcinomas yet not in adult hepatocytes (Figure 3A-C). These included Afp, encoding the fetal serum protein alfa feto-protein (Camp et al., 2017; Kuhlmann, 1978; Ruoslahti & Seppälä, 1979); Spp1, encoding osteopontin (Abu El Makarem et al., 2011; Cabiati et al., 2017; Wang et al., 2016) and Cdh17, encoding a cadherin protein associated with activation of Wnt signaling in hepatocellular carcinomas (Bartolomé et al., 2014; L. X. Liu et al., 2009) (Figure 3C). Consistently, interface hepatocytes exhibited elevated levels of Wnt pathway target genes such as Lgr5 and Axin2 (Zhao et al., 2019) (Figure 3A-B). We used smFISH to validate the specific expression of these genes (Figure 3C), as well as of Apoa1 and Actb (Figure 3D) in the hepatocytes residing at the interface between the damaged and non-damaged zones.

**Figure 3.**
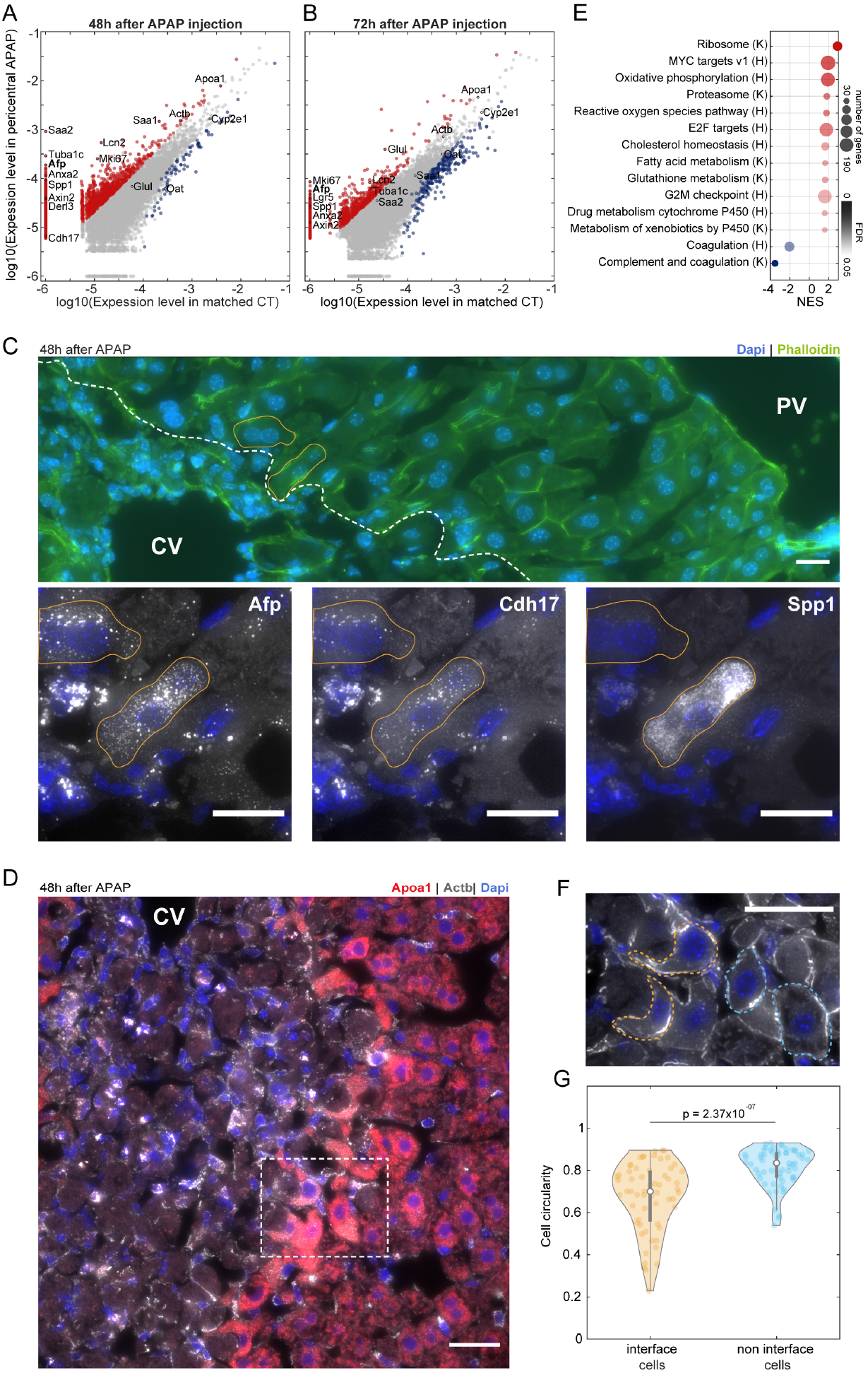
Interface hepatocytes upregulate onco-fetal genes as they reprogram into pericentral hepatocytes. (A-B) Expression levels of genes in pericentral hepatocytes 48h (A) or 72h (B) after APAP injection plotted against their expression in control hepatocytes with matched distribution of lobule spatial coordinates. Grey dots represent all genes. Red/blue dots represent genes upregulated/downregulated respectively in the regenerating tissue, with mean expression level of above 5×10^−6^, at least 2-fold difference from matched control and FDR threshold of 0.01. (C) smFISH (top) and insets (bottom) of a liver lobule 48h after APAP injection. Dashed white line delineate the damage border. CV – central vein, PV – portal vein. Nuclei and membranes are labeled with Dapi (blue) and phalloidin (green) respectively. Two representative interface hepatocytes outlined in orange shown in the insets (bottom), together with the smFISH labeling for mRNAs of AFP (left), Cdh17 (middle) and Spp1 (right). Scale bar - 20*μm*. Laplacian of gaussian filter was applied on the smFISH images. (D) smFISH of a liver lobule 48h-post APAP injection for Apoa1 (red) and Actb (grey) mRNA. Nuclei are stained with Dapi (blue). CV – central vein. Scale bar - 20*μm*. White dashed rectangle is the region displayed in (F). (E) GSEA normalized enrichment scores (NES) of gene pathways significantly upregulated (red) or downregulated (blue) in interface cells. Dot size corresponds to number of pathway genes found in the dataset; opacity corresponds to false discovery rate (FDR). Genes sets used for the analysis were taken from either KEGG (K) or Hallmark (H). (F) Magnification of the region marked in (D) showing interface (orange dashed lines) and non-interface (blue dashed lines) hepatocytes. Nuclei and membranes are labeled with Dapi (blue) and phalloidin (grey), respectively. Scale bar - 20*μm*. (G) Quantification of circularity of interface (orange) and non-interface (blue) hepatocytes. White dots represent group circularity median. Grey boxes mark the 25-75 percentiles. Significance level was calculated using paired signrank test (n=60 pairs of interface and adjacent non-interface hepatocytes, taken from 3 mice 48h after APAP injection and 3 mice 72h after APAP injection).

Interface hepatocytes upregulated pathways that included ribosomes and proteasomes (Figure 3E, Figure S2). These programs may serve to translate new pericentral proteins, while actively degrading the existing periportal/mid-lobule proteins of these migrating hepatocytes. We found that oxidative phosphorylation was also up-regulated in interface hepatocytes, in line with the elevated ATP requirements of active translation (Flamholz et al., 2014). Additionally, pericentral programs such as xenobiotic metabolism (Halpern et al., 2017) were up-regulated, whereas periportal programs such as complement and coagulation (Halpern et al., 2017) were down-regulated. Interface hepatocytes exhibited a mesenchymal-like shape with elongated extrusions extending into the damage zone (Figure 3D, F), with lower circularity compared to non-interface hepatocytes (paired ranksum pval = 2.37×10^−7^, Figure 3G). These results suggest that interface hepatocytes do not simply switch from a periportal/mid-lobular state to a pericentral state as they are pushed into the pericentral zone. Rather, their change in cellular identity is associated with a transient expression of fetal genes, elevation of protein translation and degradation and modified cellular morphology.

### Hepatic Stellate Cells exhibit spatial division of labor

Successful liver regeneration requires tightly coordinated responses from all liver cell types. HSCs are key players in liver damage response and regeneration (Friedman, 2008; Puche et al., 2013). Upon damage sensing, retinol storing quiescent HSCs are activated in a TGFb-dependent manner and transdifferentiate into collagen-secreting myofibroblasts (Baricos et al., 1999; Tipton & Dabbous, 1998). These activated stellate cells have a central role in extracellular matrix remodeling and fibrosis (Friedman, 2008; Kisseleva & Brenner, 2006; Puche et al., 2013). Previous studies used scRNAseq to characterize the acute phase of HSC activation (Kolodziejczyk et al., 2020), identifying key cytokines that may be involved in myeloid recruitment. We sought to explore the zonal dynamics of HSCs throughout the regeneration process.

We re-clustered the HSC single cells in our atlas and integrated it with a previous scRNAseq dataset (Kolodziejczyk et al., 2020) of HSCs in control and 20h post APAP (Methods, Figure 4A). HSCs have been shown to exhibit zonated gene expression along the lobule axis (Dobie et al., 2019), however the massive changes in HSC gene expression upon activation may modulate the zonal patterns of HSC landmark genes. To extract zonated HSC landmark genes, we therefore used smFISH to validate a periportal abundance of the HSC landmark gene Ngfr (Dobie et al., 2019) across all time points (Figure S3). We next selected a panel of landmark genes with high single-cell correlations with Ngfr at each time point and used these to classify HSCs into pericentral, mid-lobular and periportal zones (Methods, Figure 4B). We grouped HSCs from each time and zone to generate spatio-temporally resolved HSC expression profiles (Table S5).

**Figure 4.**
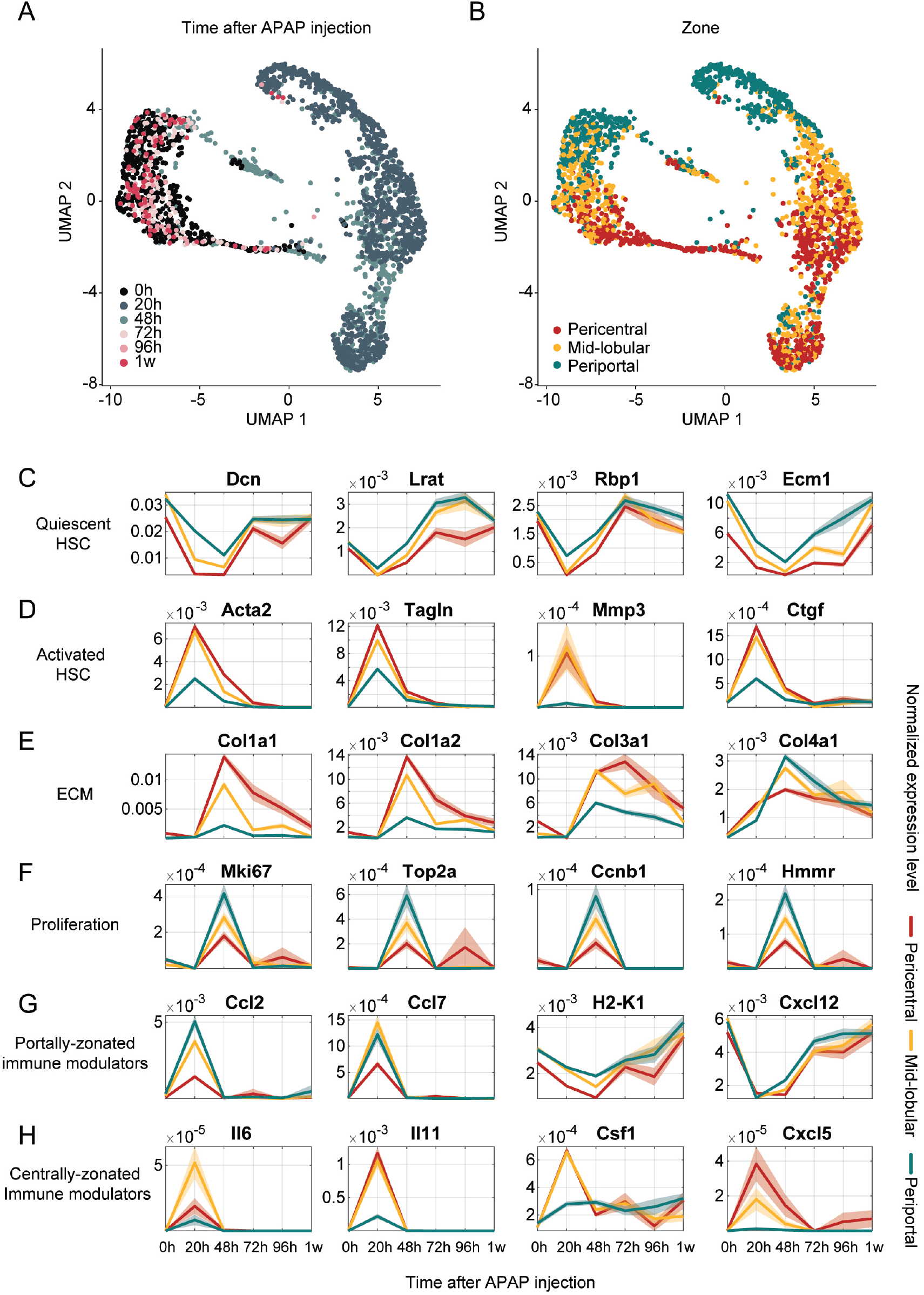
Hepatic stellate cells exhibit distinct expression programs in different zones. (A) UMAP visualization of HSCs, colored by time after APAP administration. n = 2,311 cells, at least 2 mice per time point. (B) UMAP visualization of HSCs, colored by inferred zone (Methods). (C-I) Temporal dynamics of selected genes in HSCs, stratified by zone. Lines denote the mean of the normalized expression over cells from the same zone and time point, patches denote the standard errors of the means (SE). For each gene, the mean and SE for pericentral HSCs (red), mid-lobule HSCs (yellow) and periportal HSCs (green) are presented. Selected genes belong to various processes: markers of quiescent HSCs (C), activated HSCs (D), collagen genes (E), proliferation (F), portally-zonated immune modulators (G), and centrally zonated immune modulators (H).

The levels of quiescent HSC genes such as the retinol binding protein Rbp1 and lecithin retinol acyltransferase (Lrat) declined at all zones and reverted to control levels at 96h (Figure 4C). Conversely, genes involved in HSC activation, such as the smooth muscle actin gene Acta2 acutely peaked in expression at 20h, mainly in pericentral HSCs (Figure 4D). ECM proteins such as Col1a1 and Col3a1 exhibited a delayed activation, peaking at 48h, again most prominently in pericentral HSCs, in line with the required matrix buildup in the damaged zone (Figure 4E). Notably, HSC proliferation genes, which peaked at 48h exhibited a higher expression in the periportal and mid-lobule zones (Figure 4F). Our measurements therefore suggests a potential spatial division of labor, whereby pericental HSCs rapidly activate and produce ECM, whereas periportal HSCs proliferate, potentially generating backup HSCs that can migrate into the damaged zone to support the active regeneration process.

In addition to their role in matrix remodeling, HSCs play an important role in the recruitment of myeloid cells into the liver (Pellicoro et al., 2014). We found that zonal HSC populations expressed distinct immune-modulatory proteins. Periportal HSC expressed Ccl2 and Ccl7, cytokines that recruit circulating monocytes (Jia et al., 2008; Shi & Pamer, 2011) (Figure 4G), whereas pericentral and mid-lobular HSCs expressed Il6, Il1, the macrophage regulator of proliferation and function, Csf1 (Chitu & Stanley, 2006) and Cxcl5 (Figure 4H). Notably, the lymphocyte chemoattractant Sdf-1, encoded by the gene Cxcl12 (Bleul et al., 1996) was strongly down-regulated by all HSCs. This is consistent with the lack of increase in lymphocyte abundance that we have observed throughout the zonal regeneration process (Table S3). Consistent with this avoidance of lymphocyte activation, HSCs at all zones also exhibited a decline in MHC-class 1 genes such as H2-k1 (Figure 4G). Our analysis highlight zone-specific HSC expression programs that may facilitate the temporally coordinated processes of immune recruitment, ECM buildup and breakdown that the liver exhibits following acute APAP damage.

### Endothelial cells exhibit zonated cues along the regeneration process

Liver endothelial cells are critical modulators of liver function. They form the building blocks of blood vessels, clear endotoxins, bacteria and other compounds, regulate host immune responses to pathogens, present antigens and secrete morphogens that regulate hepatocyte zonal gene expression patterns (Poisson et al., 2017). Importantly, similarly to hepatocytes and HSCs, endothelial cells exhibit zonated expression programs (Halpern et al., 2018; Inverso et al., 2021; Xiong et al., 2019). We next asked how zonal endothelial sub-populations support the regeneration process. Our atlas included 6,527 endothelial cells (Figure 5A). We used a previously established panel of zonated endothelial landmark genes such as the pericentral Wnt2 and the periportal Efnb2 (Halpern et al., 2018) (Figure 5B), as well as markers for sinusoidal endothelial cells (Kit, Halpern et al., 2018) and vascular endothelial cells (Vwf, Kalucka et al., 2020) to classify the endothelial cells into 5 zonal population (Methods). These included pericentral liver vascular endothelial cells (PC-LVEC), pericentral, mid-lobular and periportal liver sinusoidal endothelial cells (PC-LSEC, mid-LSEC and PP-LSEC) and periportal liver vascular endothelial cells (PP-LVEC, Figure 5C, Methods).

**Figure 5.**
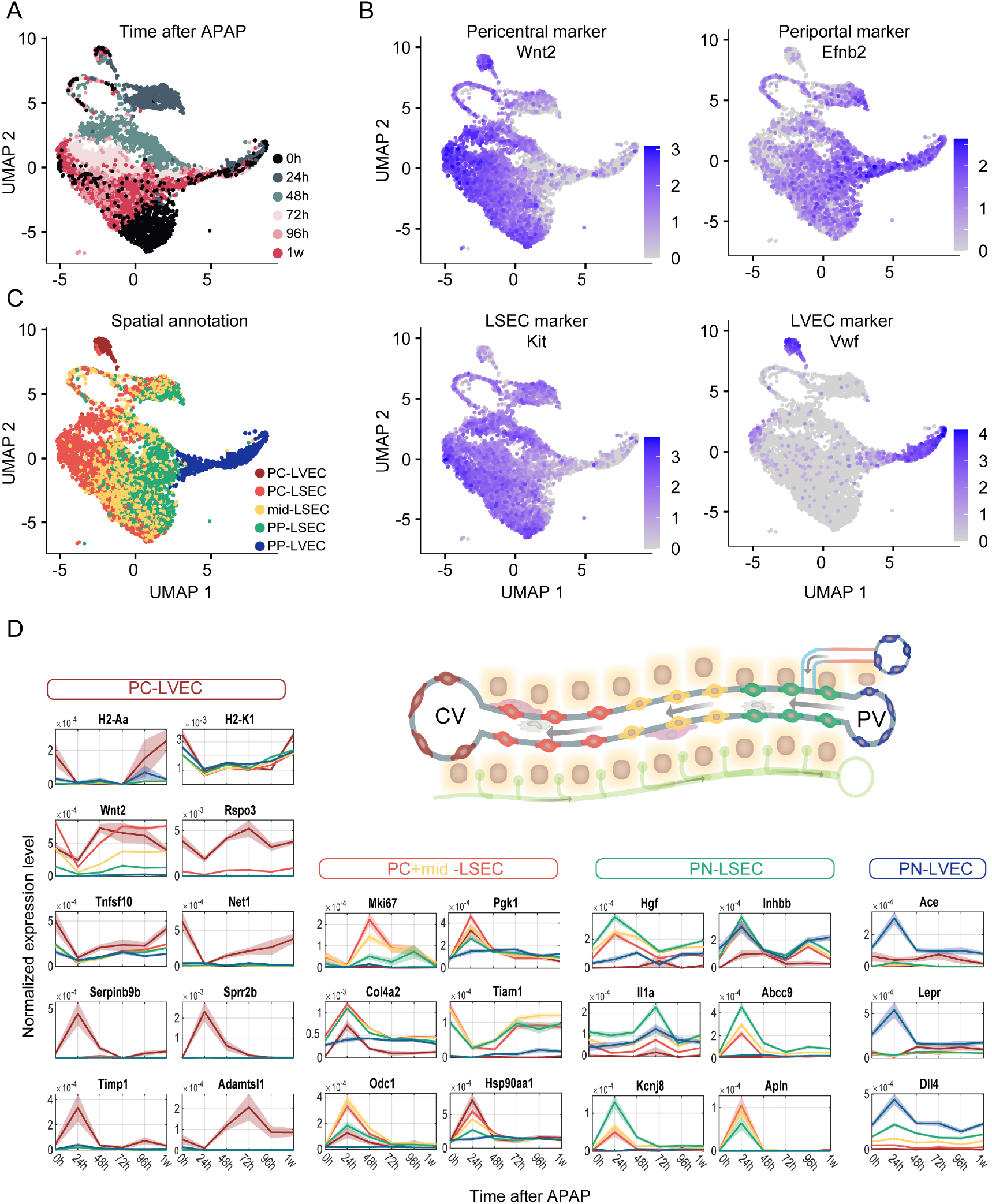
Zonal endothelial cell populations differentially express genes involved in the regeneration process. (A) UMAP visualization of endothelial cells, colored by time after APAP administration. n = 6,527 from 13 mice, at least 2 mice per time point. (B) UMAP visualization of endothelial cells, colored by expression level of the pericentral Wnt2 (top left), periportal Efnb2 (top right), sinusoidal endothelial cell marker Kit (bottom left) and vascular endothelial cell marker Vwf (bottom right). (C) UMAP visualization of endothelial cells, colored by inferred zone (Methods). PC-LVEC – pericentral liver vascular endothelial cells, PC-LSEC – pericentral liver sinusoidal endothelial cells, mid-LSEC – mid-lobule liver sinusoidal endothelial cells, PP-LSEC – periportal liver sinusoidal endothelial cells, PP-LVEC – periportal liver vascular endothelial cells. (D) Temporal dynamics of selected genes in endothelial cells, stratified by the zonated endothelial cell populations. Lines denote the mean of the normalized expression over cells from the same zone and time point, patches denote the standard errors of the means (SE). For each gene, the mean and SE for PC-LVEC (dark red), PC-LSEC (red), mid-LSEC (yellow), PP-LSEC (green) and PP-LVEC (blue) are presented. Lobule diagram highlights the different zonated endothelial cell subtypes with their respective colors.

Pericentral vascular endothelial cells, a major source of the morphogens Wnt2 and Rspo3 that establish many of the hepatocyte pericentral programs (Benhamouche et al., 2006; Mak & Png, 2020; Monga, 2015; Planas-Paz et al., 2016) showed a transient decline in the expression of these morphogens at 24h and an ensuing slight overshoot (Figure 5D). PC-LVEC also exhibited a decline in MHC-class 1 and MHC-class 2 molecules. This is in line with the decline we observed in HSC MHC-1 expression (Figure 4G), and could serve to avoid unwanted lymphocyte activation by the presentation of antigens scavenged from necrotic hepatocytes. PC-LVEC expressed the metalloproteinase inhibitor Timp1 at 24h and the metalloproteinase Adamtsl1 at 72h, consistent with the need to counteract ECM breakdown at the initial stages of regeneration and apply ECM breakdown during resolution. Pericentral and mid-lobular sinusoidal endothelial cells showed the most notable up-regulation of proliferation genes such as Mki67 (Figure 5D). This could indicate that, in addition to hepatocytes, pericentral LSECs at the damage zone are also dying and replaced by proliferation of remaining neighboring cells. Hgf, a major mitogen for hepatocytes (Nakamura et al., 1984), was broadly expressed by sinusoidal, yet not vascular endothelial cells, with a slight periportal zonation bias (Figure 5D). This broad expression could account for the broad pattern of hepatocyte proliferation we have observed (Figure 2K, L). Periportal vascular endothelial cells showed an early activation of genes such as Ace, Lepr and Dll4 at 20h. In summary, endothelial cells exhibit zonated patterns of proliferation, antigen presentation, matrix remodeling and immune recruitment throughout the regeneration process.

### Dynamics of myeloid cell subtypes along the regeneration process

Myeloid cells are instrumental to the regeneration process following acute liver damage (Goldin et al., 1996; Kolodziejczyk et al., 2020; Tacke & Zimmermann, 2014). Our atlas included 9,338 myeloid cells, clustered into 10 cell types with distinct marker (Figure 6A-C, Figure S4). Monocytes peaked in proportion at 24h whereas macrophages peaked at 48h, potentially indicating maturation from incoming monocytes, as previously shown in the liver after damage (Figure 6A,C, Karlmark et al., 2009; Zigmond et al., 2014). Kupffer cells exhibited a distinct activated cell state at 24h and 48h, yet returned to their control states by 96h (Figure 6A-C). Importantly, Kupffer cells and monocytes formed distinct clusters at all time-points (Ginhoux & Guilliams, 2016). This is consistent with the previous observation that dying Kupffer cells are replaced via proliferation of surviving Kupffer cells, rather than from incoming monocytes after acute liver damage (Zigmond et al., 2014). We analyzed the temporal expression programs of Kupffer cells, macrophages and monocytes, identifying distinct signals that may be relevant to the regeneration process (Figure S4B). As with HSCs and endothelial cells, all three myeloid cell types exhibited a drop in expression of genes encoding the antigen presentation machinery, such as Cd74 and H2-Q4 (Figure S4B). Consistently, lymphocyte-attracting chemokines such as Cxcl9, Cxcl10 and Cxcl13 were also downregulated (Figure S4B).

**Figure 6.**
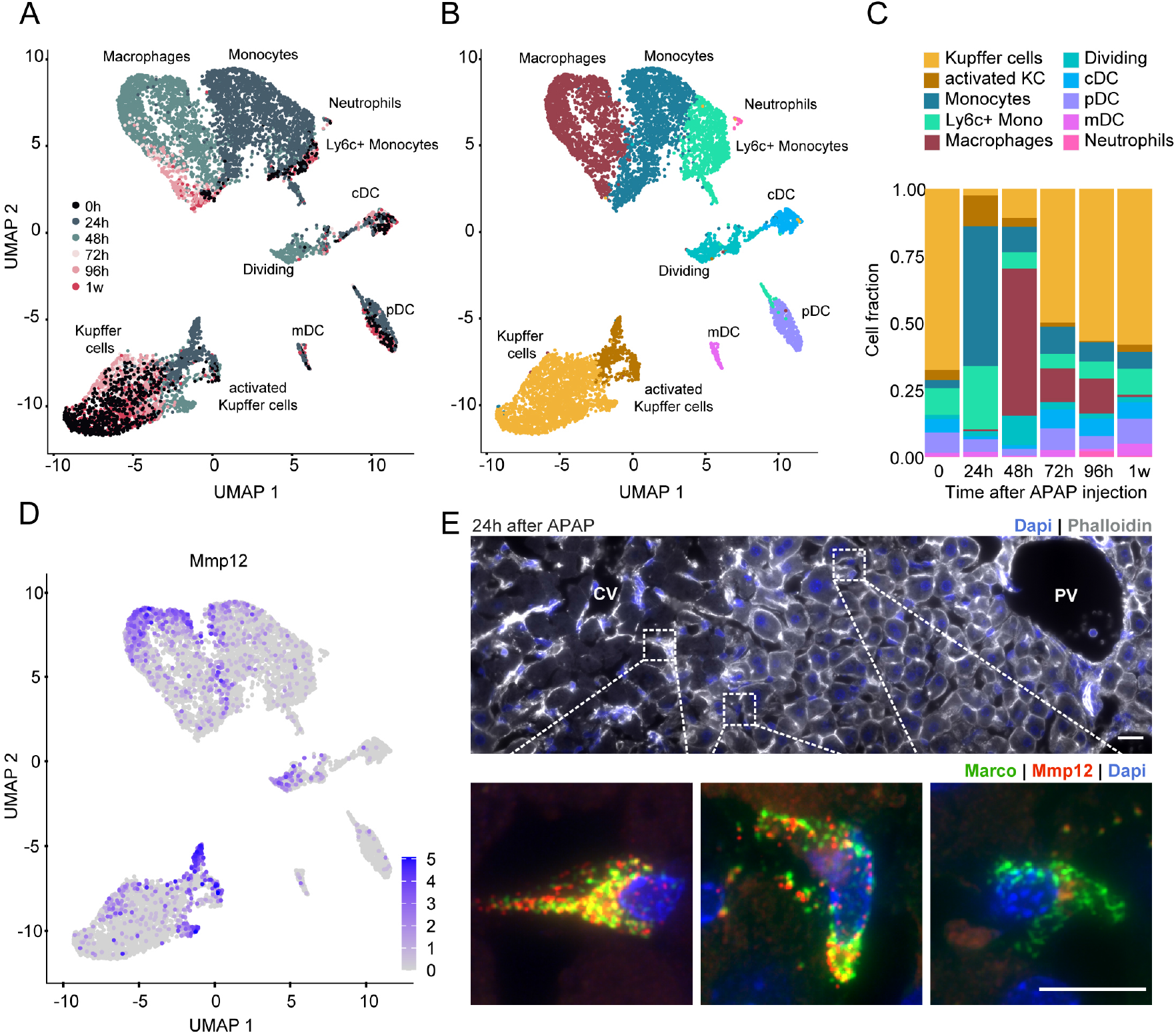
Spatio-temporal patterns of myeloid cell gene expression. (A) UMAP visualization of myeloid cell populations, colored by time after APAP administration. n = 9,338 cells from 15 mice, at least 2 mice per time point. (B) UMAP visualization of myeloid cell populations, colored by cell types. (C) The fractions of different myeloid cell subtypes at each time point. (D) UMAP visualization of myeloid cells, colored by expression levels of Mmp12. (E) smFISH scan (top) and zoomed-in insets (bottom) of a liver lobule 24h after APAP injection. CV – central vein, PV – portal vein. Nuclei and membranes are labeled with Dapi (blue) and phalloidin (green), respectively. Pericentral (left), mid-lobule (middle) and periportal (right) KCs marked by dashed lines in the scan are enlarged in the insets (bottom), together with the smFISH for the Kupffer cell gene Marco (green) and Mmp12 (red) mRNAs, together with Dapi (blue). Scale bar - 20*μm* for top image, 10*μm* for insets. Laplacian of gaussian filter was applied on the smFISH images.

The most up-regulated gene in activated Kupffer cells was Mmp12 (636.74-fold higher at 24h compared to controls, Kruskal-Wallis p (df_5,2838_) =3.37×10^−127^, Table S2), encoding for the elastase matrix metallopeptidase 12 enzyme (Figure 6D-E, Belaaouaj et al., 1995; Werb & Gordon, 1975). Mmp12 also exhibited up-regulation in macrophages (Figure 6D). Notably, when imaging the expression of Mmp12 using smFISH, we found it to be expressed in myeloid cells localized at the damage zone (Figure 6E). Both Mmp12+ Kupffer cells and MMP12+ macrophages upregulated genes found in lipid-associated macrophages (Jaitin et al., 2019; Remmerie et al., 2020), such as Trem2, Lpl and Cd36 (Figure S4B).

## Discussion

Liver zonal regeneration is a remarkable process achieved rapidly and precisely by the coordinated action of multiple cell types. These cell types exhibit both zone-independent and zone-specific gene expression programs (Figure 7). Hepatocyte proliferation occurs throughout the lobule axis, presumably to generate a large mitotic pressure that will seal the damaged zone rapidly. This broad proliferation could be induced by hepatocyte growth factor, which we found to be expressed by HSCs and sinusoidal endothelial cells at all zones, rather than exclusively at the pericentral damaged zone. We found that the incoming mid-lobular hepatocytes do not directly trans-differentiate into pericentral hepatocytes. Rather, interface hepatocytes that are pushed pericentrally to seal the damaged zone exhibit a distinct gene expression signature not observed in adult livers. This signature includes genes such as Afp, Cdh17 and Spp1, which are normally expressed in the fetal liver or in hepato-cellular carcinoma. Afp has been shown to be expressed at the margins of necrotic tissues following CCl_4_ intoxication (Iwai et al., 2000), as well as around fibrotic bridges in chronic liver damage (Nakano et al., 2017). Moreover, serum levels of the AFP protein following acute APAP damage correlate with survival (Schiødt et al., 2006; Schmidt & Dalhoff, 2005; Singh et al., 2019), consistent with our detected robust expression in reprogramming zonal hepatocytes. We found that interface hepatocytes have a distinct non-circular morphology and elevated levels of ribosomes and proteasomes. This elevation of the protein turnover machinery could serve to facilitate the required rapid turnover of their proteome, as periportal proteins need to be eliminated and pericentral proteins formed.

**Figure 7.**
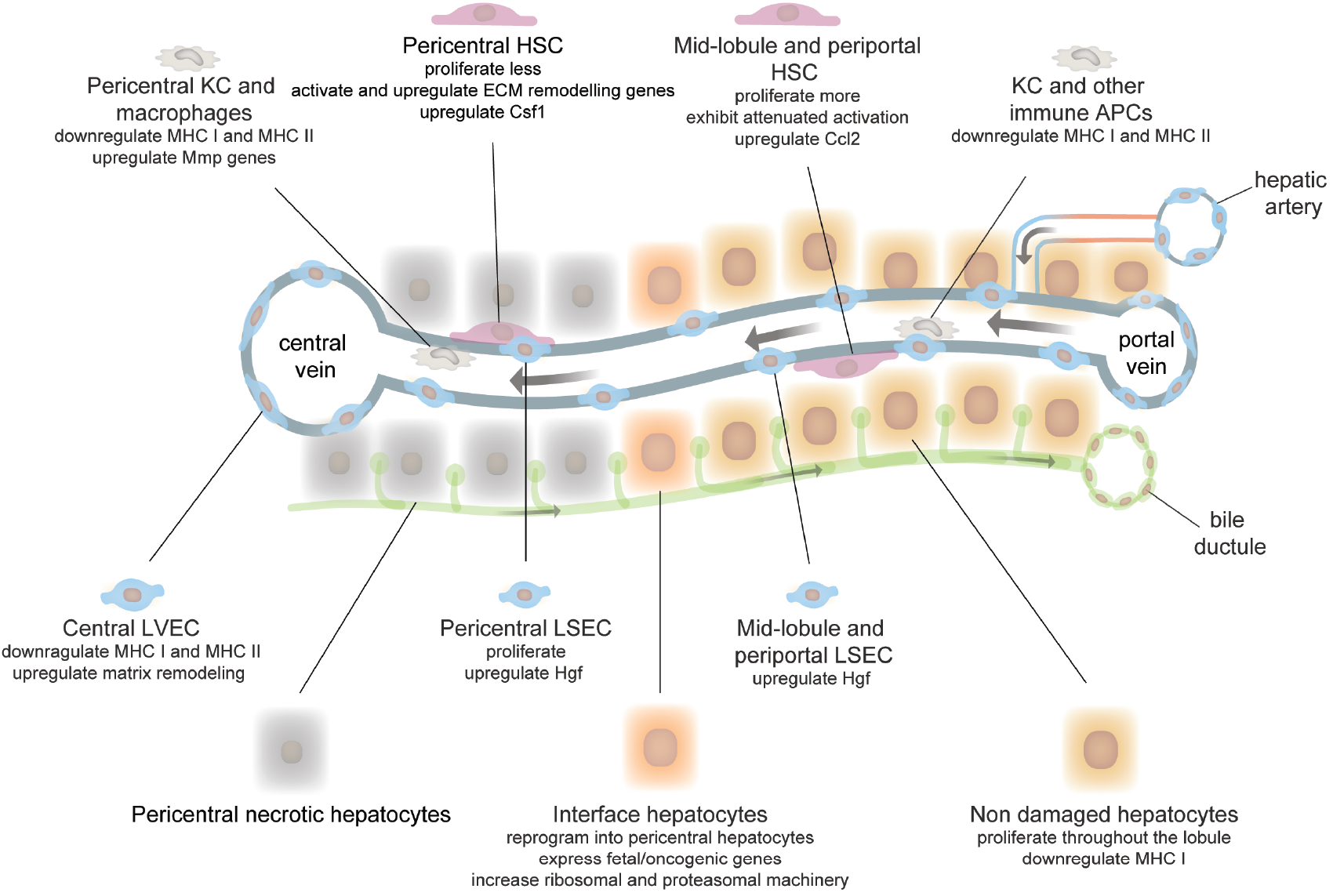
Summary of main events during APAP-induced liver damage and regeneration. A commented illustration of a liver lobule summarizing the main zone-dependent and independent changes in the different liver cell types during the regeneration process.

The fact that hepatocytes were proliferating throughout the lobule, coupled with the distinctly different zonal patterns of the pericentral Wnt morphogens and the broad Hgf mitogens, suggest that the processes of cell division and reprogramming may be uncoupled during zonal liver regeneration. The logic of this is that reprogramming needs to be performed mainly in the damaged zone, whereas broad proliferation serves to seal the damage rapidly through elevated mitotic pressure. It would be important to identify the cues that give rise to the fetal reprogramming of interface hepatocytes, be they mechanosensing of the contact with the fibrotic zones or distinct morphogens.

While zonal regeneration involved the massive transient recruitment of monocytes and macrophages, lymphocyte abundance showed stability in cell numbers (Table S3). The avoidance of induction of adaptive immunity may be critical, given the multiple new antigens available for presentation due to the necrotic hepatocytes. Indeed, we found that all cell types transiently reduce the expression of MHC-I genes and that the liver antigen presenting cells – endothelial cells and myeloid cells transiently reduce MHC-II genes up to 96h after damage. The induction in myeloid-recruiting cytokines such as Ccl2, Ccl7, Csf1, and the repression of signals for recruitment of lymphocytes, such as Cxcl12 and Cxcl13 may also contribute to the inverse abundances of myeloid and lymphocyte populations.

Our study exposed intricate zone-dependent expression programs of HSCs and endothelial cells. We found that pericentral HSCs massively induced ECM programs, such as collagen gene expression, whereas periportal HSCs showed increased proliferation as well as higher expression of Ccl2. Response to zonal damage requires massive matrix remodeling. To achieve this, the liver needs to generate new HSCs and to activate them to achieve maximal secretion of matrix components as fast as possible. Both HSC proliferation and the production of secreted matrix proteins require cellular resources that may be limiting. Our analyses suggest a strategy of spatial resource allocation, whereby pericentral HSCs are early responders that initiate matrix production immediately, whereas portal HSCs divide to provide backup cells that can later migrate into the damaged zone and reduce the load from the pericentral HSCs. Our study also provide the molecular details of the resolution of the fibrogenesis involved in zonal regeneration. These could prove informative for the goals of resolving fibrotic states (Adler et al., 2020; Henderson & Iredale, 2007; Kisseleva & Brenner, 2006; X. Liu et al., 2020; Puche et al., 2013; Troeger et al., 2012). Our analysis also revealed zonated proliferation patterns of endothelial cells, which were restricted to pericentral and mid-lobular sinusoidal endothelial cells but not pericentral vascular endothelial cells. These vascular endothelial cells, which form the central blood vessels and are non-fenestrated might be more robust to the toxic microenvironment generated following hepatocyte necrosis and myeloid cell migration.

The short time frame of four days at which the complete zonal regeneration process is attained is within the order of magnitude of typical protein lifetimes (Schwanhäusser et al., 2011). In such out-of-steady state regime a complete functional view of the regeneration process will require analysis of proteins. It will therefore be informative to apply approaches such as spatial sorting (Ben-Moshe et al., 2019; Inverso et al., 2021) to assess the dynamics of liver proteome in a cell and zone-specific manner. Our study focused on pericentral zonal damage. It will be interesting to use similar methods to explore whether interface hepatocytes, immune evasion and matrix remodeling also apply to periportal damage (Ben-Moshe & Itzkovitz, 2019). In summary, our work highlighted the details of liver plasticity during zonal regeneration and exposed the zone-dependent coordinated programs that facilitate precise healing of the liver tissue.

## Methods

### APAP injections

Mouse experiments were approved by the Institutional Animal Care and Use Committee of the Weizmann Institute of Science and were conducted in agreement with the institute guidelines. Male (C57BL/6JOlaHsd) mice aged 8-11weeks, housed under regular 12h light-dark cycle, were used in our experiments. Food (Teklad Global 18% Protein Rodent Diet) was taken out 12h prior to APAP injection. Mice had access to drinking water. A dose of 300mg/1kg body weight of APAP (Merck, 103-90-2) was injected i.p at a concentration of 40mg/ml, diluted in saline 0.9%. APAP solution was slightly heated to improve drug solubility. In order to avoid circadian rhythm effects, all timepoints were carried out at the same hour of the day – food was taken out at ZT16 (10pm) and APAP or saline was injected at ZT4 (10am). Food was given back to mice after injection mice were fed ad libitum until sacrificed for experiment.

For bulk RNA sequencing and smFISH experiments, 8 control mice were injected with saline alone following 12h fasting – at 6h, 24h, 96h and 1m, 2 mice per time point. Mice were injected with APAP 6h, 24h, 48h, 72h, 96h, 1week and 1month prior to their sacrifice, 3 mice per time point. For scRNAseq experiments, non-injected mice were used as controls (n=2) and mice were injected with APAP 24h (n=2), 48h (n=4), 72h (n=2), 96h (n=3) and 1week (n=2) prior to their sacrifice.

### Single molecule Fluorescence in situ Hybridization (smFISH)

APAP- or saline-treated mice were sacrificed 6h, 24h, 48h, 72h, 96h, 1w and 1m after injection. Mice were sacrificed by cervical dislocation. A part of the median lobe was taken for bulk mRNA sequencing, and the rest of the liver was fixed with pre-chilled 4% PFA for 3h in 4°C followed by an overnight incubation with 4% PFA and 30% sucrose in 4°C. Fixed liver was then embedded in OCT. smFISH experiments were performed on 8µm thick cryosections mounted on poly-L-lysine pre-coated coverslips. Probe libraries were designed using the Stellaris FISH Probe Designer Software (Biosearch Technologies, Inc., Petaluma, CA). Hybridization was performed according to published protocol (Lyubimova et al., 2013). Briefly, tissues were permeabilized for 10min with proteinase K (10µg/ml Ambion AM2546) followed by 2 washes of 2× SSC (Ambion AM9765) for 5min. Tissues were incubated in wash buffer (20% Formamide Ambion AM9342, 2× SSC) for 5min and then with hybridization buffer (10% Dextran sulfate Sigma D8906, 20% Formamide, 1mg/ml E.coli tRNA Sigma R1753, 2× SSC, 0.02% BSA Ambion AM2616, 2 mM Vanadylribonucleoside complex NEB S1402S) mixed with probes. Hybridization mix was incubated with tissues overnight in a 30°C incubator. smFISH probe libraries were coupled to Cy5, TMR or Alexa594. After the hybridization, tissues were washed with wash buffer containing 50ng/ml DAPI for 30 min at 30°C. DAPI (Sigma-Aldrich, D9542) was used for nuclear staining. Membranes were labeled using actin filaments staining by Phalloidin (Thermofisher). All images were performed on a Nikon-Ti-E inverted fluorescence microscope using the NIS element software AR 5.11.01. Scans spanning from central to portal veins were acquired using 100x magnification. Stitching of the individual fields was done by NIS element program, with 15% overlap.

### Bulk liver mRNA isolation and mcSCRBseq library construction

Mice injected with APAP or saline vehicle only were sacrificed using cervical dislocation. A small part from the median liver lobe, between 2-5mm^3^, was taken for bulk RNA sequencing, and the rest of the organ was fixed for smFISH experiments, as aforementioned. Tissue samples were placed in cold 600µl TRI-reagent for RNA isolation (Sigma). 0.5mm diameter RNase free zirconium-oxide homogenization beads (Next Advance) were added in a mass comparable to that of the tissue sample, and samples were homogenized in a Bullet Blender (Next Advance) using speed 8 setting for 3mins. After the homogenization step, samples were centrifuged (14,000 rpm, 30sec) and 500ul from the supernatant were transferred into a new Eppendorf tube. An equal volume of 100% EtOH was added to the sample. From that tube, 500µl of sample were transferred into Direct-zol RNA miniprep column (Zymo research). RNA extraction was performed as detailed in kit protocol, with a DNase I incubation step. Total RNA was eluted in 20µl nuclease free water and 1µl of total RNA was taken for library construction. Libraries were prepared according to mcSCRBseq protocol (Bagnoli et al., 2018). Following reverse transcription and exo-nuclease steps, cDNA was pre-amplified with 10-15 cycles, depending on cDNA concentration, indicated by qPCR quality control. 2ng of the amplified cDNA were used to library construction, using Nextera XT DNA Library kit (Illumina, FC-131-1024), according to manufacturer protocol. Quality control of the resulting libraries was performed with an Agilent High Sensitivity D1000 ScreenTape System (Agilent, 5067-5584). Libraries were loaded with a concentration of 2.2pM on 75 cycle high output flow cells (Illumina, FC-404-2005) and sequenced on a NextSeq 500/550 (Illumina) with the following cycle distribution: 8bp index1, 16 bp read1, 66 bp read2 (no index2 needed), with estimated depth of 15M reads per sample. A total of 29 libraries for 29 different mice were sequenced.

### Data processing for bulk mRNA

Illumina output files were demultiplexed with bcl2fastq v.2.17. Resulting FASTQ files were analyzed using the zUMIs pipeline (Parekh et al., 2018). Reads were aligned with STAR (v.2.5.3a) to the GRCm38 genome (release 84; Ensembl) and exonic unique molecular identifier (UMI) counts were used for downstream analysis. Two samples (CT_24h_m9 and CT_24h_m10) were discarded for having a total UMI sum of less than 1.5M UMIs. Data was further normalized by dividing each sample by its sum of UMIs, resulting in normalized expression levels corresponding to the gene’s fraction in the sample.

### Principal component analysis on bulk samples

Fractions were natural-log transformed (after the addition of the minimal fraction larger than zero) and Z-scores of the log transformed fractions of each gene were calculated across all samples. PCA was calculated on the Z-scores. In order to avoid circadian effects, control mice which were injected with saline 6h prior to the experiment were discarded from the pool of control mice.

### Liver perfusions and hepatocyte dissociation

Mice were anaesthetized with 100 mg/kg Ketamine (Zoetis Manufacturing & Research) and 10 mg/kg Xylazine (EuroVet) dissolved in 1× PBS and injected i.p. Once anaesthetized, their livers were perfused as previously described (Mederacke et al., 2015), with some adjustments. A 27 G syringe connected to perfusion line and pump was inserted into the vena cava; 7-10 ml of pre-warmed to 37 °C EGTA buffer were perfused to wash the blood from the liver. After EGTA, 12-20 ml of pre-warmed to 37 °C enzyme buffer solution (EBS) with 2.3 U of Liberase Blendzyme 3 recombinant collagenase (Roche) were cannulated into the vena cava to isolate hepatocytes. Shortly after the beginning of the perfusion, the portal vein was cut to allow drainage of the blood. Successfully perfused livers were extracted to a Petri dish with 25 ml of pre-warmed EBS and gently minced using forceps. Dissociated liver cells were collected and filtered through a 100µm cell strainer. Cells were spun down at 30g for 3 min at 4 °C to obtain the hepatocyte-enriched pellet. Pellet was resuspended in 25ml cold EBS. To enrich for live hepatocytes, 22.5 ml Percoll (Sigma-Aldrich) mixed with 2.5 ml 10× PBS was added to the cells. Cells were centrifuged at 600 rpm for 10min at RT. The supernatant containing the dead cells was aspirated and cells were resuspended in Cell Staining Buffer (BioLegend).

### Liver perfusions of non-parenchymal cell (NPC) dissociation

Perfusion procedure for NPC dissociation was similar to that of hepatocytes, although it required the use of different dissociating enzymes. Instead of using Liberase Blendzyme 3, livers were sequentially perfused with 7-10ml 37°C EGTA, then 10-15 ml of pre-warmed to 37 °C enzyme buffer solution (EBS) containing 0.4 mg/ml Protease (Roche) and then with 15-20 ml of 37 °C EBS containing collagenase D (0.1 U/ml) (Roche). Damaged livers (48h-72h after APAP injection) were perfused with 1.5x concentration of dissociating enzymes. Livers were extracted to a Petri dish with 25 ml of pre-warmed EBS and gently minced using forceps. Dissociated liver cells were filtered through a 70µm cell strainer. Cells were spun down at 30g for 3 min at 4 °C twice, and each time the pellet was discarded to eliminate the hepatocytes. Supernatant was then centrifuged at 580g for 10min in 4°C. Pellet containing NPCs was resuspended in 1ml Red Blood Cell Lysis Buffer (Sigma), incubated at room temp for 1min at RT. EBS was then added and samples were centrifuged again in 580g for 10min in 4°C. Pellet was resuspended in 100-200µl Cell Staining Buffer (Biolegend).

### TotalSeq™-B for 10x Feature Barcoding

We multiplexed our samples using a hashing technique (Stoeckius et al., 2018), by labeling samples with oligonucleotide-barcoded antibodies. To this end, we used commercial tagged antibodies - TotalSeq™-B for 10x Feature Barcoding (bioLegend), according to manufacturer protocol. A small fraction of the cells was stained with 50% trypan blue to validate that dead cells do not make up more than 25% of the sample. Hepatocytes / NPCs were counted and 2M cells in a volume of 100µl Cell Staining Buffer were taken for further preparation. Cells from each mouse were blocked with 5µl of FCX (BioLegend) for 10min in 4°C. While cells were incubating in blocking solution, Total seq B antibodies (BioLegend) were prepared using 1µg of each TotalSeq™ hashtag antibody. Unique antibodies were then added to each sample and were incubated for 30min in 4°C. To wash out the antibodies, samples were filled to 10ml with Cell Staining Buffer and centrifuged 500g in 4°C for 5 minutes 3 times. Finally, cells were counted and cells viability was estimated using trypan blue, to make sure no more than 30% cells of the samples are dead.

### Single Cell Gene Expression with Feature Barcoding technology

Two to four antibody-tagged hashed samples were multiplexed into one well of 10x Chromium Next GEM Chip G, loading approximately 10,000 cells per well (2,500-5,000 cells per sample). Single cells were processed using the 10x Chromium Next GEM Single Cell 3’ Reagent Kits v3.1 with Feature Barcoding technology for Cell Surface Protein, according manufacturer’s manual. Libraries were sequenced using NovaSeq 6000 using NovaSeq Sp (100 cycles) sequencing Kits (Illumina). A total of 7 sequencing runs of cells taken from 19 mice: 15 mice for NPC (2 non injected control mice, 2mice 20h after APAP, 4mice 48h after APAP, 2mice 72h after APAP, 3mice 96h after APAP and 2mice 1w after APAP), and 4 mice for hepatocytes (2mice for 48h after APAP and 2mice for 72h after APAP).

### Mapping Single-cell RNAseq data analysis

Single-cell RNA-seq data were demultiplexed, aligned to the GRCm38 mouse genome assembly (mm10) and UMIs were quantified using the Cell Ranger Single-Cell Software Suite 3.1.0 and bcl2fastq, with the functions “cellranger mkfastq” and “cellranger count”.

### Data pre-processing and background subtraction for all cells

Starting from the raw matrix output of Cell Ranger, droplets with low counts of UMIs were filtered out. Threshold was set to 1,000 UMIs, however, in sequencing runs with a 99.9th quantile of sum of UMIs of below 1000, threshold was reduced to 500 UMIs. Of the remaining droplets, sum of UMIs for each droplet were log10-transformed, and Otsu’s method (Otsu, 1979) was used to determine a threshold that separates empty from cell-containing droplets (Macosko et al., 2015). Cells were defined as droplets with log10 UMI sums above this threshold. Background was defined as droplets with log10 UMI sums below the threshold. These droplets potentially contained floating RNA from dead or ruptured cells. The mean UMI count of each gene across the background droplets was calculated. This vector was subtracted from each cell. The same averaging and subtraction were performed for UMI counts of the feature barcodes. Negative values were corrected to zero. This background subtraction was done independently for each sequencing run.

### Data processing of NPCs using Seurat package

Background-subtracted cells’ gene expression matrices were further analyzed using the R package Seurat 3.1.5 (Stuart et al., 2019). Cells with UMI sums of above 3,000 UMIs and mitochondrial gene percent less than 15% were retained. Only cells with maximal fraction of feature barcode above 70% were included, to avoid doublets. Genes detected in a minimum of 5 cells were retained. Data were normalized and scaled using the SCTransform function, with regression of sum of UMIs (vars.to.regress = “nCount_RNA”). PCA was calculated based on the variable genes found in the previous function, however, mitochondrial (“^mt-”) and ribosomal (“^Rp[ls]”) genes were manually removed from the gene list since they are prone to batch-related expression variability. Following PCA, number of PCs for clustering was determined using the elbow plot method (between 15-20 PCs). Clustering was performed with default parameters. Resulting clusters were then manually annotated after extracting cluster markers using FindAllMarkers Seurat function. Clusters containing more than 50% cells with expression of markers of more than one cell type (for example cluster of cells expressing both HSC marker Dcn and Kupffer cell marker Clec4f) were removed and remaining cells were retransformed, re-clustered and re-annotated.

### Data integration of NPCs

Seurat objects from all sequencing runs were integrated using the following Seurat functions: ‘SelectIntegrationFeatures’, with mitochondrial and ribosomal genes excluded from the output 5,000 genes; ‘PrepSCTIntegration’ with default settings; and ‘FindIntegrationAnchors’ and ‘IntegrateData’ with ‘SCT’ as the normalization method. PCA and clustering on integrated Seurat object was done as detailed above for the individual Seurat objects. Clusters were annotated to the main liver resident / infiltrating immune cell types: endothelial cells, HSC, hepatocytes, cholangiocytes, Kupffer cells (KC), monocytes, macrophages, plasmacytoid dendritic cells (pDC), conventional dendritic cells (cDC), B cells, T cells and natural killer (NK) cells.

### Hepatocyte Processing

Our experimental data included hepatocytes from two mice sacrificed at 48 hours and two mice at 72 hours after APAP injection, dissociated using Liberase enzyme. In order to include control non-injected hepatocytes, hepatocytes from a previously published study (Droin et al., 2021) were integrated. Notably, the animals were held at the same facility prior to experiment as animals used for our experiments and cells isolation and library construction were performed similarly to the APAP experiments. From their data set, the two mice perfused at a similar circadian time as our mice (samples ‘ZT6A’ and ‘ZT6B’) were chosen. Fatsq files of these two mice were re-aligned and preprocessed as detailed above for our sequencing runs, to eliminate any bioinformatics biases. Individual Seurat objects were created for the two mice, filtering out cells with less than 1,000 total UMI count and with mitochondrial percentage of less than 9% and more than 35%, as was found to capture viable hepatocytes (Droin et al., 2021). These included 923 cells for ZT6A and 1,076 cells for ZT6B. Next, we created a Seurat object of our background-removed APAP-injected hepatocytes to include cells with UMI count of at least 2,500 UMIs and mitochondrial percent of 9%-35% with fraction of maximal feature barcode above 70%. Doublets were removed from the dataset after clustering, as described above for NPCs. The Seurat object included 532 hepatocytes from 2 mice 48h after APAP injection, and 239 hepatocytes from 2 mice 72h after APAP injection.

### Data integration of Hepatocytes

Integration of control mice and APAP injected mice was performed in the same manner as for NPCs. In total, 2770 cells from either control, 48h or 72h after APAP injection time points were used for hepatocyte analysis. Integrated hepatocyte cluster was then integrated with NPC clusters, using the same integration parameters described above.

### Finding markers for the different cell types

To plot the heatmap of differentially expressed genes between the different cell types (Figure S1A), a signature matrix containing the mean normalized expression of the genes in each of the 11 cell type clusters was computed. The matrix included only genes with maximal normalized mean expression above 5×10^−4^. Our markers for each cell type included genes for which the expression was highest for the respective cell and more than 2-fold higher than any other cell type. We further retained for each cell type, the 8 genes with the highest ratio as markers.

### Estimating abundance of each cell type from bulk liver RNAseq dataset using CIBERSORTx

To avoid biases originating in dissociation and sensitivity differences between the different cell types, we imputed the abundance of each of the 11 main cell types found in our scRNAseq dataset in each of the time points after APAP injection using CIBERSORTx computional deconvolution software (Newman et al., 2019). To this end, the “impute cell fractions” module of the online tool was used, with the bulk liver RNAseq dataset of 17,972 genes expressed in 29 samples as the mixture file. The mean expression per cell type in our single cell data was used for the signature matrix. First, expression of each cell was normalized to the cell’s sum of UMI counts, resulting in expression fractions. Next, the mean expression of genes was calculated for each cell type. The resulting matrix of 19,344 genes across 11 cell types was the input for the signature matrix file. CIBERSORTx was executed with the default parameters.

### Extracting hepatocyte gene expression temporal changes from bulk liver RNAseq data

To follow the temporal changes of hepatocytes throughout the liver regeneration time course (Figure 2A), bulk liver RNAseq data was used. First, hepatocyte specific genes were selected by detecting genes with mean expression level of at least 10^−4^ and 20-fold higher in the hepatocyte cluster in the scRNAseq dataset than any of the other 10 cell type clusters. Mup genes were excluded from the list of genes, given their high variability. Next, 328 out of the resulting 342 hepatocyte specific genes were found in the bulk RNAseq dataset. Each gene was further normalized by dividing the sum of the 328 genes in each sample. Spearman correlation was then calculated between the normalized expression of these genes for all pairs of four control samples (not including 6h control due to circadian differences and 24h controls, due to insufficient coverage) and APAP-treated samples.

### Inferring lobule spatial coordinates of single hepatocytes

Several studies have established algorithms for computationally assigning dissociated single cells into their tissue coordinates, based on known spatially differential gene expression (Achim et al., 2015; Droin et al., 2021; Halpern et al., 2017, 2018; Moor et al., 2018). Expression of known zonated genes, termed landmark genes, provides spatial information about the origin of the hepatocyte, along the one-dimensional porto-central axis. We implemented the same method described in Droin et al., 2021 (Droin et al., 2021). In short, 19 pericentrally zonated (Alad, Aldh1a1, C6, Cpox, Csad, Cyp1a2, Cyp2c37, Cyp2c50, Cyp2e1, Cyp3a11, Gstm1, Hpd, Lect2, Mgst1, Oat, Pon1, Prodh, Rgn, Slc16a10) and 31 periportally zonated (Apof, Apom, Asgr2, Ass1, Atp5a1, C8b, Cpt2, Cyp2f2, Eef1b2, Elovl2, Fads1, Fbp1, Gc, Gnmt, Hsd17b13, Ifitm3, Igf1, Igfals, Ndufb10, Pigr, S100a1, Serpina1c, Serpina1e, Serpind1, Serpinf1, Trf, Uqcrh, Vtn, Pck1, Arg1, Cps1) landmark genes were used. The gene lists are comprised of genes selected by Droin and colleagues (Droin et al., 2021) which appeared in our dataset. These genes underwent two-step normalization: first, the genes were divided by the sum of total UMI count of each cell, to reflect the fraction of expression of each landmark gene in each cell. Second, the genes were scaled by dividing them by their maximal expression across all cells. This provided similar weight to all landmark genes, regardless of expression levels differences among them. Then, for each cell, we summed up the normalized-scaled levels of all central landmark genes (cLM) and all portal landmark genes (pLM). For each cell, we then calculated the zonation coordinate as pLM ⁄(cLM+pLM). The resulting quotient corresponded to a lobule coordinate from pericentral to periportal, respectively. To assign each cell to three discrete zones – pericentral, mid-lobule or periportal, for each time point, we sorted the zonation coordinates. Cells with the lowest third zonation coordinates were ascribed to pericentral zone, the highest third of cells was ascribed to periportal zone, and the middle third was ascribed to mid-lobule zone. To extract zonation profile of genes at different time points, the means and standard errors of the means were calculated for cells from same time point and zone.

### Analyzing proliferation in hepatocytes

To analyze the proliferation at different zones and time points, we used the cell cycle genes provided in Seurat package (‘cc.genes$s.genes’ and ‘cc.genes$g2m.genes’). We found 94 genes present in our dataset from the list of 97 cell-cycle related genes, and calculated their summed SCT-normalized expression levels in each of the cells.

### Comparing interface hepatocytes and control

In order to systematically define pericentral hepatocytes in the regenerating tissues according to the zonation coordinate, the median of the zonation coordinates in control mice hepatocytes was calculated. Central hepatocytes in 48h and 72h after APAP injections were defined as cells with zonation coordinate lower than the median control zonation coordinate (0.73). Consequently, zonation coordinate of pericentral hepatocytes ranged between 0.45 - 0.72 for 48h time point and 0.31 - 0.72 for 72h time point. To further compare the expression signatures of these cells with control pericentral genes from same lobule layers, 100 control hepatocytes were sampled to match the distribution of zonation coordinates of the APAP-injected pericentral hepatocytes using the inverse transform sampling method (Devroye, 2006).

### Gene Set Enrichment Analysis (GSEA) of interface hepatocytes gene expression

To identify gene pathways enriched in interface hepatocytes, Spearman correlations of genes with the expression of Afp, which was found to be specifically expressed in interface cells, were calculated. Cells included in this analysis were only from 48h and 72h after APAP injection and from pericentral and mid-lobule zones, where interface hepatocytes were detected. Next, the top 99th percentile genes with the highest correlation with Afp (108 genes) from each of the two time points were summed up and the Spearman correlation of individual genes with this meta-signature was calculated. Included in this analysis were genes, the mean sum-normalized expression of which was higher than 5×10^−6^ in at least one time point. The list of the remaining genes was ranked by their Spearman correlation and was used as input in GSEA tool, for preranked genes (Mootha et al., 2003; Subramanian et al., 2005). Gene list was analyzed against KEGG and Hallmark gene sets datasets.

### Measuring nucleus diameter of regenerated hepatocytes

H&E images of control, 1 week after APAP and 1month after APAP were scanned. In total 1,518 hepatocytes were analyzed. Pericentral and periportal populations were defined as the two adjacent hepatocyte layers surrounding the vein. The nuclear diameter was measured by Case Viewer software. The diameters of the nuclei were matched to ploidy levels sizes was based on previous study (Tanami et al., 2017).

### Hepatic stellate cell cluster analysis

The integrated cluster of HSCs included 465 cells from 4 mice 48h after APAP injection, 98 cells from 2 mice 72h, 31 cells from 96h and 78 cells from 1w after APAP injection. However, our sequencing experiments failed to capture sufficient numbers of cells from control and 24h after APAP injection. To complement the dataset, we used a recently published dataset (Kolodziejczyk et al., 2020). Again, the animals were held at the same facility prior to experiment as animals used for our experiments and cells isolation and library construction were done in a similar manner to ours. Nonetheless, in this study, APAP injection was given to non-fasted mice in a concentration of 500mg/Kg and cells were sorted prior to loading on 10x chromium. We used seven wt mice: three non-treated mice (‘SFP_1’, ‘SPF_2’, and ‘SPF_3’ mice) and four mice injected with APAP 20h prior to perfusion (‘SFP_APAP_1’, ‘SPF_ APAP_2’, ‘SPF_ APAP_3’, and ‘SPF_ APAP_2’). Fatsq files of these seven mice were re-aligned and preprocessed as detailed above for our sequencing runs, to eliminate any bioinformatics biases. Individual Seurat objects were created for the seven mice, filtering out cell with less than 3,000 total UMI count and with mitochondrial percentage of more than 15%. The Seurat objects were then integrated as described above and clusters of HSC (annotated by markers found using ‘FindAllMarkers’ function) were subset. These included 563 cells from 3 control mice and 1,073 cells from 4 mice 20h after APAP injection. Next, we integrated HSC from our experiments and from Kolodziejczyk et al. using same integration method described above. The few cells from 24h after APAP from our measurements were pooled with cells from 20h after APAP. Overall, HSC cluster included 2,311 HSCs from six pooled time points.

### Inferring lobule spatial coordinates of single HSCs

Zonation of quiescent and activated HSCs was previously studied (Dobie et al., 2019). Ngfr was found to be highly correlated with periportal and pericentral zonation, respectively (Dobie et al., 2019). Given the notable transcriptional changes induced by liver damage and HSC activation, smFISH was used to validate that Ngfr was consistently expressed at higher levels in periportal genes. Ngfr was therefore used as our main periportal landmark gene. Due to the low capture rate and dropouts associated with scRNAseq, we sought to increase the number of landmark genes for each time point. To this end, single-cell Spearman correlations of each gene with Ngfr were calculated, for each time point separately. Expression levels were normalized to the sum of UMI count for each cell. Only genes with mean expression levels over all cells in the specific time point higher than 10^−4^ and correlation higher than 0.25 were considered for first iteration. This yielded a signature of six periportal genes appearing in at least five out of the six time points: Ngfr, Plvap, Colec11, Sod3, Steap4 and Ifitm1. In a second iteration, the correlation of each gene with the summed expression level of these six periportal landmark genes was calculated. For each time point, genes with mean expression higher than 10^−4^ and correlation higher than 0.4 or lower than -0.4 and correlation pval lower than 0.01 were considered as landmark genes – positively correlated genes as portal landmark genes and negatively correlated genes as central landmark genes. Mitochondrial (‘^mt-‘) and ribosomal (‘^Rp[ls]’) genes were excluded from all landmark gene lists. After establishing the portal and central landmark genes for each time point, further analysis and zone assignment were performed as described above for hepatocytes.

### Endothelial cell cluster analysis

Integrated cluster of endothelial cells was comprised of 6,527 cells, 978 cells from 2 control mice, 654 cells from 2 mice 24h after APAP injection, 1,234 cells from 2 mice 48h after APAP injection, 1,229 cells from 2 mice 72h after APAP injection, 797 cells from 3mice 96h after APAP injection and 1,635 cells from 2 mice 1 week after APAP injection.

### Inferring lobule spatial coordinates of single endothelial cells

Spatial variability of endothelial cells is originating from both the location along the lobule porto-central and from the endothelial cell subtype – either liver vascular endothelial cells (LVECs), or liver sinusoidal endothelial cells (LSEC) (Kalucka et al., 2020). Endothelial cells were thus divided into five zonated populations: pericentral (PC-) LVEC, PC-LSEC, mid-lobule (mid) LSEC, periportal (PP-) LSEC and PP-LVEC. The endothelial cells were first subset into a separate Seurat object, reclustered using Seurat. Clusters expressing the vascular endothelial cell marker Vwf (Kalucka et al., 2020) were annotated as LVEC clusters. Of those, the cluster expressing both Vwf and Wnt2, a prominent pericentral marker (Halpern et al., 2018; Wang et al., 2015), was annotated as PC-LVEC and the remaining cluster was annotated as PP-LVEC. Out of the remaining cells (LSECs), Zonation reconstruction was implemented as described above for hepatocytes. The landmark genes selected for reconstruction were taken from previous study (Halpern et al., 2018).

### Myeloid clusters analysis

The integrated NPC dataset included 9,338 cells in five main myeloid clusters: Kupffer cells, monocytes, macrophages, pDCs and cDCs. These cells were subset to a separate Seurat object, and cells were processed and clustered as detailed for the individual sequencing runs. 10 PCs were used for clustering, and resolution was set to 0.2. Markers for each of the resulting 10 clusters were found using ‘FindAllMarkers’ function, and clusters were subsequently annotated to different cell types and cell states, using ImmGen database (available at http://www.immgen.org/) and previous studies (Deczkowska et al., 2021; Kolodziejczyk et al., 2020).

## Supporting information

Table_S1_bulkRNA_seq_UMI_table

Table_S2_single_cell_data_mean_SEM_by_cell_type_per_time_point

Table_S3_CIBERSORTx_Results

Table_S4_heaptocyte_data_by_tp_and_zone

Table_S5_HSC_data_by_tp_and_zone

Table_S6_Endo_data_by_tp_and_zone

## Data availability

Data generated in this study available at the Zenodo repository under the following URL: https://doi.org/10.5281/zenodo.5172137

## Code availability

All code used will be available from the authors upon request.

## Acknowledgments

S.I. is supported by the Wolfson Family Charitable Trust, the Edmond de Rothschild Foundations, the Fannie Sherr Fund, the Dr. Beth Rom-Rymer Stem Cell Research Fund, the Minerva Stiftung grant, the Israel Science Foundation grant No. 1486/16, the Broad Institute-Israel Science Foundation grant No. 2615/18, the European Research Council (ERC) under the European Union’s Horizon 2020 research and innovation program grant No. 768956, the Chan Zuckerberg Initiative grant No. CZF2019-002434, the Bert L. and N. Kuggie Vallee Foundation and the Howard Hughes Medical Institute (HHMI) international research scholar award. We thank Ehud Zigmond from Tel-Aviv Sourasky Medical Center for fruitful discussions and Merav Kedmi, Orit Zion and Hadas Keren-Shaul from Weizmann Institute Life Sciences Core Facilities for their assistance in sequencing.

## Supplementary Figures

**Figure S1 (related to Figure 1) –.**
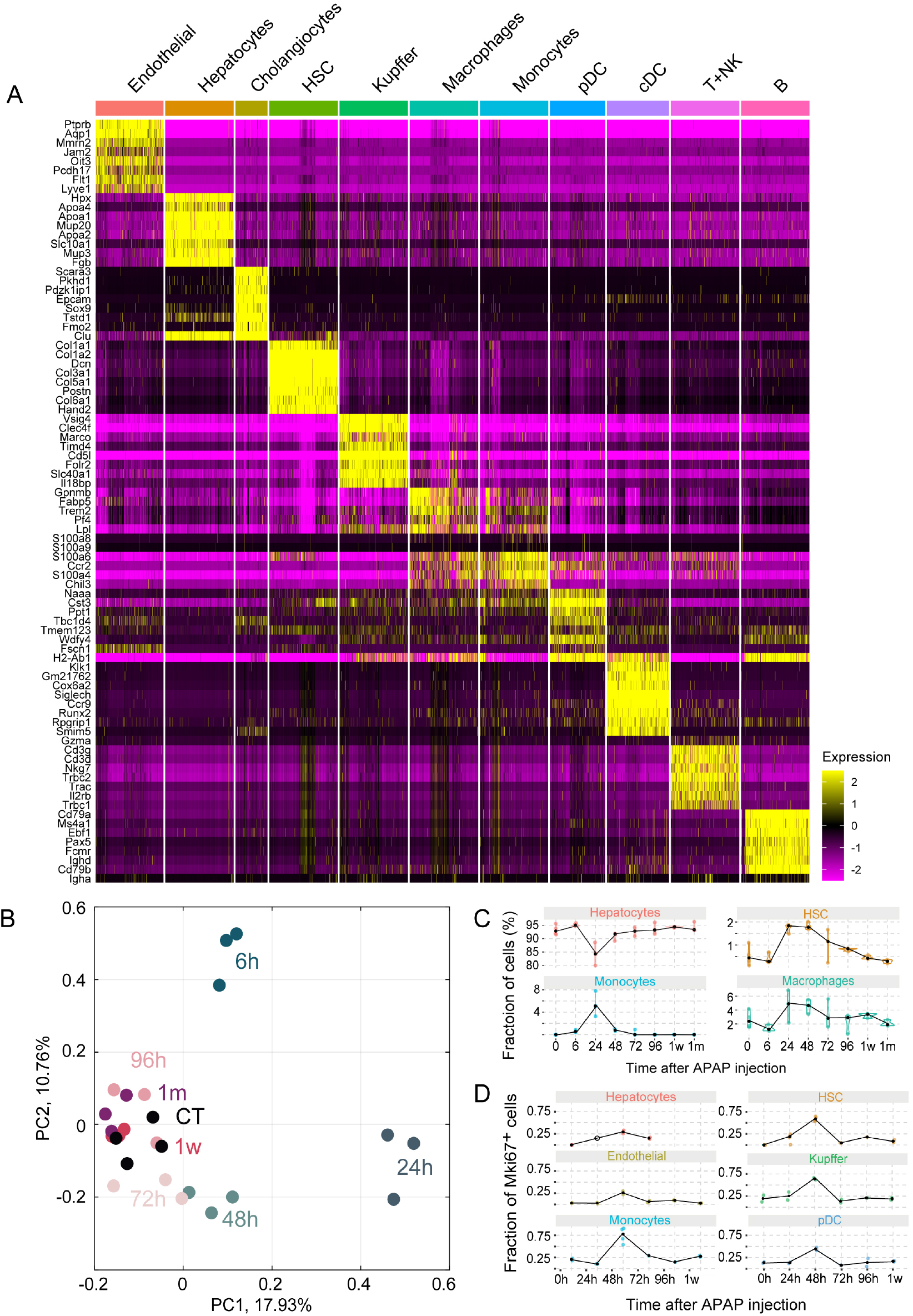
Cell type markers and distributions along the regeneration time-course. (A) Heatmap of the expression of markers for each cell type (Methods). Expression is SCTransformed, cells randomly down-sampled to a maximum of 500 per cell type. (B) Principal component analysis of the sequenced bulk liver samples. The two first components are plotted. Samples are colored by the time following the APAP injection. (C) The estimated fractions of selected cell types over time. Each dot is a mouse. Black dots mark the medians. The fractions were estimated using CIBERSORTx computational deconvolution of the bulk measurements, using the mean single cell data as cell-type signature (Methods). (D) Fractions of single cells that are positive for the proliferation marker Mki67. Each dot is a mouse. Only mice with a minimum 20 cells per cell type per time point were considered in this analysis. Black dots marks the median of all mice. White dot denotes missing data (hepatocytes at 24h).

**Figure S2 (related to Figure 3) –.**
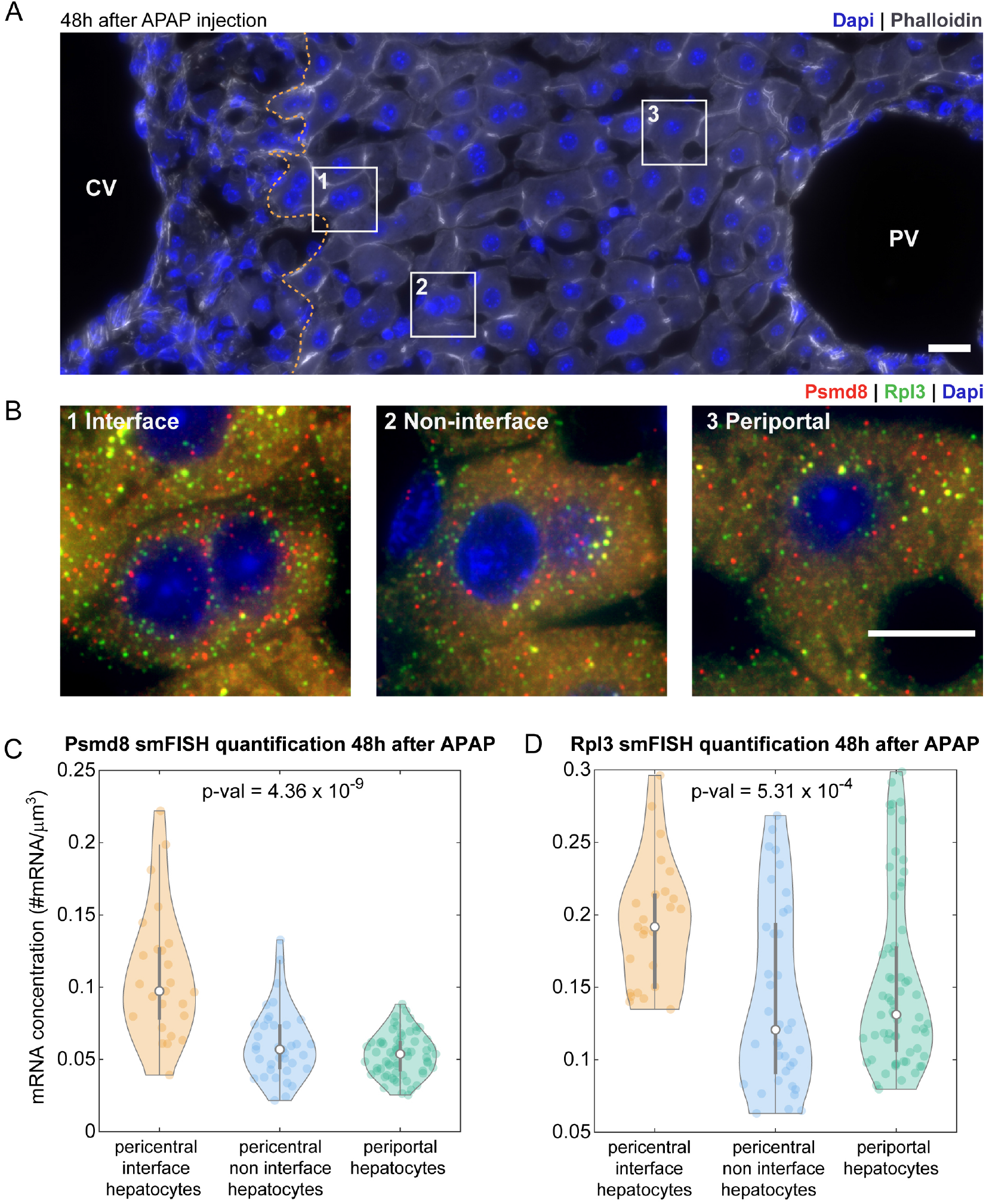
Interface hepatocytes upregulate ribosomal and proteasomal genes. (A-B) smFISH (A) and insets (B) of a liver lobule 48h after APAP injection. CV – central vein, PV – portal vein. Nuclei and membranes are labeled with Dapi (blue) and Phalloidin (grey), respectively. Orange dashed line demarcates border of damaged area. Shown in insets smFISH images of Rpl3 (green) and Psmd8 (red) mRNA in interface pericentral (1), non-interface pericentral (2), and periportal (3) hepatocytes 48h after APAP injection. Scale bars - 20*μm* (A), 10*μm* (B). Laplacian of gaussian filter was applied on the smFISH images. (C-D) smFISH quantification of Psmd8 (C) and Rpl3 (D) transcripts in mice 48h following APAP injection. n = 125 cells from 2 mice. P-value was calculated using Kruskal Wallis test (df_2,124_).

**Figure S3 (related to Figure 4) –.**
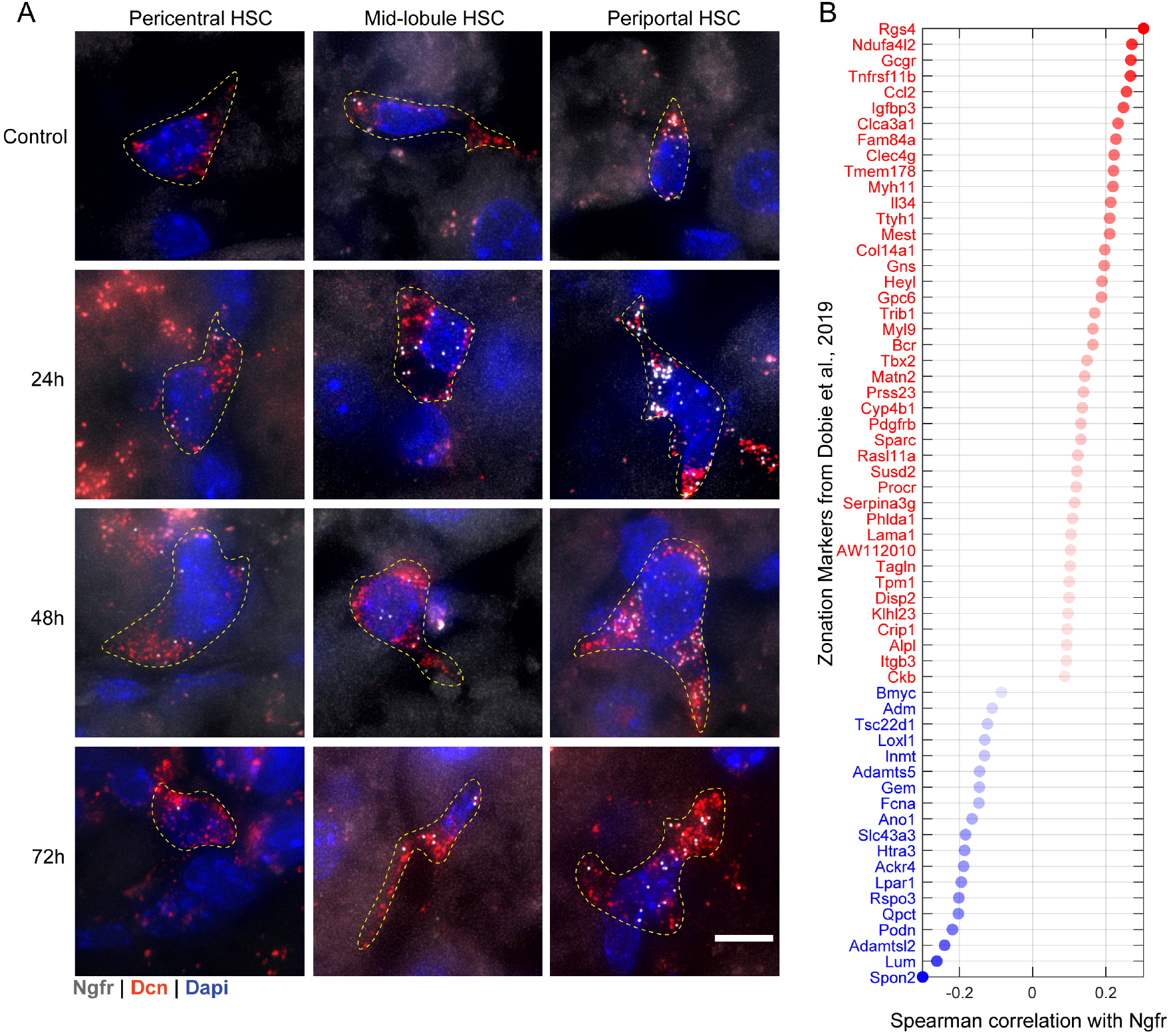
Ngfr is periportally zonated throughout the regeneration process and can serve as a landmark gene. (A) smFISH images of pericentral (left), mid-lobule (middle) and periportal (right) HSCs at different time points after APAP injection. Dashed lines demarcate HSC contours. HSCs were identified by Dcn mRNA expression (red dots). Grey dots – single transcripts of Ngfr. Dapi labeled nuclei are blue. Scale bar - 10*μm*. (B) Spearman correlations of pericentrally (blue) and periportally (red) zonated gene expressions with Ngfr expression level in control single HSCs. List of zonated genes was taken from previously published study by Dobie et al., 2019, only genes with significant correlation (p-val < 0.05) are presented. Dot opacity corresponds to p-val.

**Figure S4 (related to Figure 6) –.**
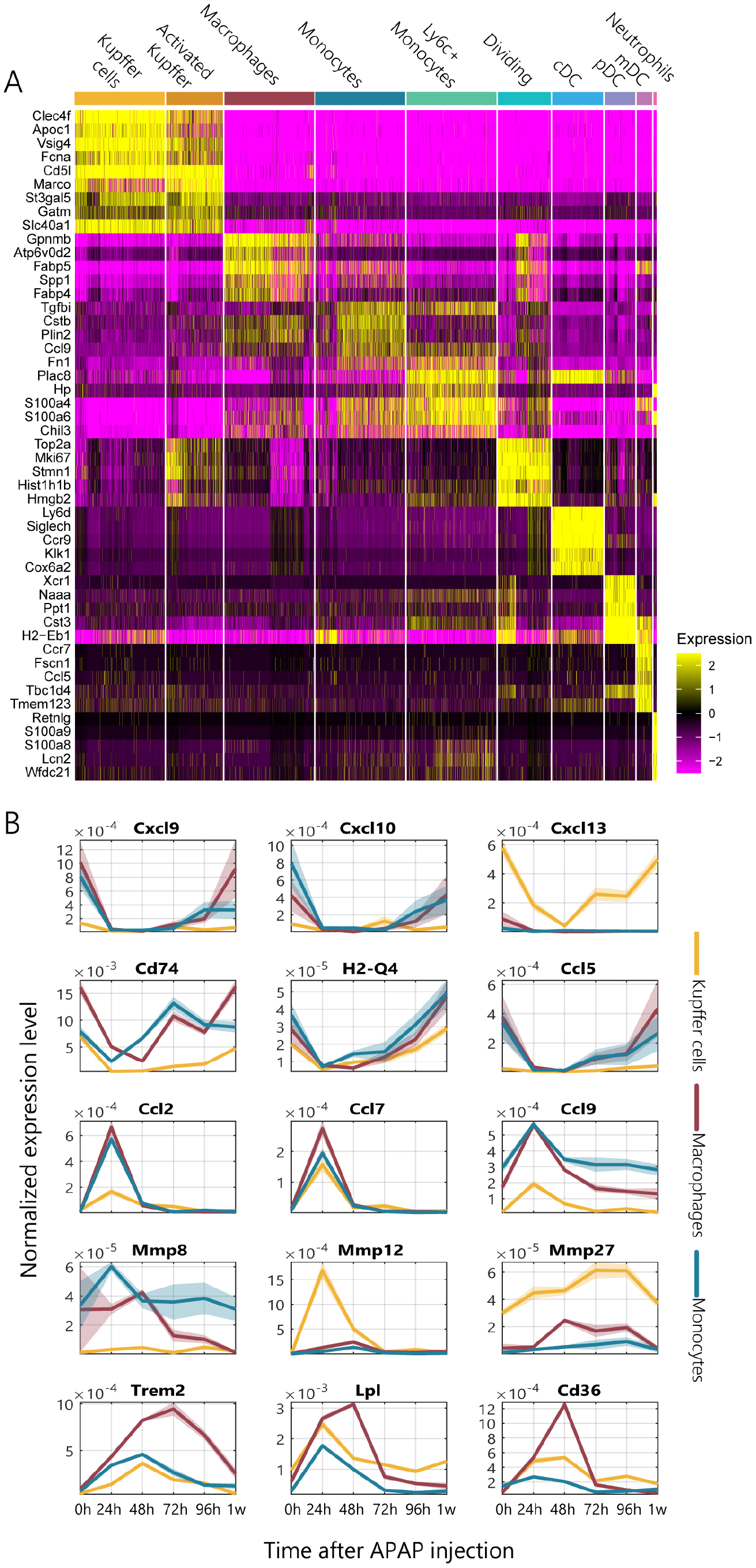
Markers of the myeloid subtypes and temporal dynamics of selected genes. (A) Heatmap of the expression of markers for each myeloid sub-type (Methods). Expression is SCTransformed, cells randomly down-sampled to a maximum of 500 per cell type. (B) Temporal dynamics of selected genes in Kupffer cells (yellow), macrophages (purple) and monocytes (blue). Lines denote the mean of the normalized expression over cells from the same time point, patches denote the standard errors of the means (SE).

## Supplementary Data

**Table S1 – bulk RNA sequencing UMI table**

UMI count matrix of bulk RNA libraries of 29 mice liver samples in seven different time points after APAP injection and saline-injected controls.

**Table S2 – temporal mean expressions of the different cell types**

Mean expression and standard error of the mean (SEM) of the eleven cell types stratified by the six time points. Mean and SEM were calculated on fractions of genes out of the sum of UMIs in each cell.

**Table S3 – computational deconvolution of bulk Liver samples into the different cell types using CIBERSORTx**

CIBERSORTx output file showing the inferred fractions of each of the main eleven cell type clusters in our scRNAseq data in the bulk liver samples (Methods).

**Table S4 - temporal mean expressions of hepatocytes stratified by zones**

Gene expression of zonally-stratified hepatocytes at different time points. Shown are means and standard errors of the means (SEM). Units are fraction of cellular UMIs.

**Table S5 - temporal mean expressions of hepatic stellate cells stratified by zones**

Gene expression of zonally-stratified hepatic stellate cells at different time points. Shown are means and standard errors of the means (SEM). Units are fraction of cellular UMIs.

**Table S6 - temporal mean expressions of endothelial cells stratified by zones**

Gene expression of zonally-stratified endothelial cells at different time points. Shown are means and standard errors of the means (SEM). Units are fraction of cellular UMIs.

## Notes

https://doi.org/10.5281/zenodo.5172137

### Competing Interest Statement

The authors have declared no competing interest.

## References

Abu El Makarem, M. A., Abdel-Aleem, A., Ali, A., Saber, R., Shatat, M., Rahem, D. A., & Sayed, D. (2011). Diagnostic significance of plasma osteopontin in hepatitis C virus-related hepatocellular carcinoma. Annals of Hepatology, 10(3), 296–305. https://doi.org/10.1016/S1665-2681(19)31541-8

Adler, M., Mayo, A., Zhou, X., Franklin, R. A., Meizlish, M. L., Medzhitov, R., Kallenberger, S. M., & Alon, U. (2020). Principles of Cell Circuits for Tissue Repair and Fibrosis. IScience, 23(2), 100841. https://doi.org/10.1016/j.isci.2020.100841

Baricos, W. H., Cortez, S. L., Deboisblanc, M., & Xin, S. (1999). Transforming Growth Factor-β Is a Potent Inhibitor of Extracellular Matrix Degradation by Cultured Human Mesangial Cells. Journal of the American Society of Nephrology, 10(4), 790–795. https://doi.org/10.1681/ASN.V104790

Bartolomé, R. A., Barderas, R., Torres, S., Fernandez-Aceñero, M. J., Mendes, M., García-Foncillas, J., Lopez-Lucendo, M., & Casal, J. I. (2014). Cadherin-17 interacts with α2β1 integrin to regulate cell proliferation and adhesion in colorectal cancer cells causing liver metastasis. Oncogene, 33(13), 1658–1669. https://doi.org/10.1038/onc.2013.117

Belaaouaj, A., Shipley, J. M., Kobayashi, D. K., Zimonjic, D. B., Popescu, N., Silverman, G. A., & Shapiro, S. D. (1995). Human Macrophage Metalloelastase. GENOMIC ORGANIZATION, CHROMOSOMAL LOCATION, GENE LINKAGE, AND TISSUE-SPECIFIC EXPRESSION *. Journal of Biological Chemistry, 270(24), 14568–14575. https://doi.org/10.1074/jbc.270.24.14568

Benhamouche, S., Decaens, T., Godard, C., Chambrey, R., Rickman, D. S., Moinard, C., Vasseur-Cognet, M., Kuo, C. J., Kahn, A., Perret, C., & Colnot, S. (2006). Apc Tumor Suppressor Gene Is the “Zonation-Keeper” of Mouse Liver. Developmental Cell, 10(6), 759–770. https://doi.org/10.1016/j.devcel.2006.03.015

Ben-Moshe, S., & Itzkovitz, S. (2019). Spatial heterogeneity in the mammalian liver. Nature Reviews Gastroenterology & Hepatology, 16(7), 395–410. https://doi.org/10.1038/s41575-019-0134-x

Ben-Moshe, S., Shapira, Y., Moor, A. E., Manco, R., Veg, T., Bahar Halpern, K., & Itzkovitz, S. (2019). Spatial sorting enables comprehensive characterization of liver zonation. Nature Metabolism, 1(9), 899–911. https://doi.org/10.1038/s42255-019-0109-9

Bhushan, B., & Apte, U. (2019). Liver Regeneration after Acetaminophen Hepatotoxicity: Mechanisms and Therapeutic Opportunities. The American Journal of Pathology, 189(4), 719– 729. https://doi.org/10.1016/j.ajpath.2018.12.006

Bleul, C. C., Fuhlbrigge, R. C., Casasnovas, J. M., Aiuti, A., & Springer, T. A. (1996). A highly efficacious lymphocyte chemoattractant, stromal cell-derived factor 1 (SDF-1). Journal of Experimental Medicine, 184(3), 1101–1109. https://doi.org/10.1084/jem.184.3.1101

Cabiati, M., Gaggini, M., Cesare, M. M., Caselli, C., De Simone, P., Filipponi, F., Basta, G., Gastaldelli, A., & Del Ry, S. (2017). Osteopontin in hepatocellular carcinoma: A possible biomarker for diagnosis and follow-up. Cytokine, 99, 59–65. https://doi.org/10.1016/j.cyto.2017.07.004

Camp, J. G., Sekine, K., Gerber, T., Loeffler-Wirth, H., Binder, H., Gac, M., Kanton, S., Kageyama, J., Damm, G., Seehofer, D., Belicova, L., Bickle, M., Barsacchi, R., Okuda, R., Yoshizawa, E., Kimura, M., Ayabe, H., Taniguchi, H., Takebe, T., & Treutlein, B. (2017). Multilineage communication regulates human liver bud development from pluripotency. Nature, 546(7659), 533–538. https://doi.org/10.1038/nature22796

Chitu, V., & Stanley, E. R. (2006). Colony-stimulating factor-1 in immunity and inflammation. Current Opinion in Immunology, 18(1), 39–48. https://doi.org/10.1016/j.coi.2005.11.006

Devroye, L. (2006). Chapter 4 Nonuniform Random Variate Generation. In S. G. Henderson & B. L. Nelson (Eds.), Handbooks in Operations Research and Management Science (Vol. 13, pp. 83– 121). Elsevier. https://doi.org/10.1016/S0927-0507(06)13004-2

Dobie, R., Wilson-Kanamori, J. R., Henderson, B. E. P., Smith, J. R., Matchett, K. P., Portman, J. R., Wallenborg, K., Picelli, S., Zagorska, A., Pendem, S. V., Hudson, T. E., Wu, M. M., Budas, G. R., Breckenridge, D. G., Harrison, E. M., Mole, D. J., Wigmore, S. J., Ramachandran, P., Ponting, C. P., … Henderson, N. C. (2019). Single-Cell Transcriptomics Uncovers Zonation of Function in the Mesenchyme during Liver Fibrosis. Cell Reports, 29(7), 1832-1847.e8. https://doi.org/10.1016/j.celrep.2019.10.024

Droin, C., Kholtei, J. E., Bahar Halpern, K., Hurni, C., Rozenberg, M., Muvkadi, S., Itzkovitz, S., & Naef, F. (2021). Space-time logic of liver gene expression at sub-lobular scale. Nature Metabolism, 3(1), 43–58. https://doi.org/10.1038/s42255-020-00323-1

Flamholz, A., Phillips, R., & Milo, R. (2014). The quantified cell. Molecular Biology of the Cell, 25(22), 3497–3500. https://doi.org/10.1091/mbc.E14-09-1347

Friedman, S. L. (2008). Hepatic Stellate Cells: Protean, Multifunctional, and Enigmatic Cells of the Liver. Physiological Reviews, 88(1), 125–172. https://doi.org/10.1152/physrev.00013.2007

Gebhardt, R. (1992). Metabolic zonation of the liver: Regulation and implications for liver function. Pharmacology & Therapeutics, 53(3), 275–354. https://doi.org/10.1016/0163-7258(92)90055-5

Ginhoux, F., & Guilliams, M. (2016). Tissue-Resident Macrophage Ontogeny and Homeostasis. Immunity, 44(3), 439–449. https://doi.org/10.1016/j.immuni.2016.02.024

Goldin, R. D., Ratnayaka, I. D., Breach, C. S., Brown, I. N., & Wickramasinghe, S. N. (1996). Role of Macrophages in Acetaminophen (paracetamol)-Induced Hepatotoxicity. The Journal of Pathology, 179(4), 432–435. https://doi.org/10.1002/(SICI)1096-9896(199608)179:4<432::AID-PATH609>3.0.CO;2-S

Halpern, K. B., Shenhav, R., Massalha, H., Toth, B., Egozi, A., Massasa, E. E., Medgalia, C., David, E., Giladi, A., Moor, A. E., Porat, Z., Amit, I., & Itzkovitz, S. (2018). Paired-cell sequencing enables spatial gene expression mapping of liver endothelial cells. Nature Biotechnology, 36(10), 962– 970. https://doi.org/10.1038/nbt.4231

Halpern, K. B., Shenhav, R., Matcovitch-Natan, O., Tóth, B., Lemze, D., Golan, M., Massasa, E. E., Baydatch, S., Landen, S., Moor, A. E., Brandis, A., Giladi, A., Stokar-Avihail, A., David, E., Amit, I., & Itzkovitz, S. (2017). Single-cell spatial reconstruction reveals global division of labour in the mammalian liver. Nature, 542(7641), 352–356. https://doi.org/10.1038/nature21065

Henderson, N. C., & Iredale, J. P. (2007). Liver fibrosis: Cellular mechanisms of progression and resolution. Clinical Science, 112(5), 265–280. https://doi.org/10.1042/CS20060242

Inverso, D., Shi, J., Lee, K. H., Jakab, M., Ben-Moshe, S., Kulkarni, S. R., Schneider, M., Wang, G., Komeili, M., Vélez, P. A., Riedel, M., Spegg, C., Ruppert, T., Schaeffer-Reiss, C., Helm, D., Singh, I., Boutros, M., Chintharlapalli, S., Heikenwalder, M., … Augustin, H. G. (2021). A spatial vascular transcriptomic, proteomic, and phosphoproteomic atlas unveils an angiocrine Tie–Wnt signaling axis in the liver. Developmental Cell, 56(11), 1677-1693.e10. https://doi.org/10.1016/j.devcel.2021.05.001

Iwai, M., Morikawa, T., Muramatsu, A., Tanaka, S., Mori, T., Harada, Y., Okanoue, T., Kashima, K., & Ishii, M. (2000). Biological Significance of AFP Expression in Liver Injury Induced by CCL4. Acta Histochemica Et Cytochemica, 33(1), 17–22. https://doi.org/10.1267/ahc.33.17

Jaitin, D. A., Adlung, L., Thaiss, C. A., Weiner, A., Li, B., Descamps, H., Lundgren, P., Bleriot, C., Liu, Z., Deczkowska, A., Keren-Shaul, H., David, E., Zmora, N., Eldar, S. M., Lubezky, N., Shibolet, O., Hill, D. A., Lazar, M. A., Colonna, M., … Amit, I. (2019). Lipid-Associated Macrophages Control Metabolic Homeostasis in a Trem2-Dependent Manner. Cell, 178(3), 686-698.e14. https://doi.org/10.1016/j.cell.2019.05.054

Jia, T., Serbina, N. V., Brandl, K., Zhong, M. X., Leiner, I. M., Charo, I. F., & Pamer, E. G. (2008). Additive Roles for MCP-1 and MCP-3 in CCR2-Mediated Recruitment of Inflammatory Monocytes during Listeria monocytogenes Infection. The Journal of Immunology, 180(10), 6846–6853. https://doi.org/10.4049/jimmunol.180.10.6846

Jungermann, K., & Keitzmann, T. (1996). Zonation of Parenchymal and Nonparenchymal Metabolism in Liver. Annual Review of Nutrition, 16(1), 179–203. https://doi.org/10.1146/annurev.nu.16.070196.001143

Kalucka, J., de Rooij, L. P. M. H., Goveia, J., Rohlenova, K., Dumas, S. J., Meta, E., Conchinha, N. V., Taverna, F., Teuwen, L.-A., Veys, K., García-Caballero, M., Khan, S., Geldhof, V., Sokol, L., Chen, R., Treps, L., Borri, M., de Zeeuw, P., Dubois, C., … Carmeliet, P. (2020). Single-Cell Transcriptome Atlas of Murine Endothelial Cells. Cell, 180(4), 764-779.e20. https://doi.org/10.1016/j.cell.2020.01.015

Karlmark, K. R., Weiskirchen, R., Zimmermann, H. W., Gassler, N., Ginhoux, F., Weber, C., Merad, M., Luedde, T., Trautwein, C., & Tacke, F. (2009). Hepatic recruitment of the inflammatory Gr1+ monocyte subset upon liver injury promotes hepatic fibrosis. Hepatology, 50(1), 261–274. https://doi.org/10.1002/hep.22950

Kisseleva, T., & Brenner, D. A. (2006). Hepatic stellate cells and the reversal of fibrosis. Journal of Gastroenterology and Hepatology, 21(3), S84–S87. https://doi.org/10.1111/j.1440-1746.2006.04584.x

Kolodziejczyk, A. A., Federici, S., Zmora, N., Mohapatra, G., Dori-Bachash, M., Hornstein, S., Leshem, A., Reuveni, D., Zigmond, E., Tobar, A., Salame, T. M., Harmelin, A., Shlomai, A., Shapiro, H., Amit, I., & Elinav, E. (2020). Acute liver failure is regulated by MYC-and microbiome-dependent programs. Nature Medicine, 26(12), 1899–1911. https://doi.org/10.1038/s41591-020-1102-2

Kuhlmann, W. D. (1978). Localization of ALPHA1-fetoprotein and DNA-synthesis in liver cell populations during experimental hepatocarcinogenesis in rats. International Journal of Cancer, 21(3), 368–380. https://doi.org/10.1002/ijc.2910210319

Liu, L. X., Lee, N. P., Chan, V. W., Xue, W., Zender, L., Zhang, C., Mao, M., Dai, H., Wang, X. L., Xu, M. Z., Lee, T. K., Ng, I. O., Chen, Y., Kung, H., Lowe, S. W., Poon, R. T. P., Wang, J. H., & Luk, J. M. (2009). Targeting cadherin-17 inactivates Wnt signaling and inhibits tumor growth in liver carcinoma. Hepatology, 50(5), 1453–1463. https://doi.org/10.1002/hep.23143

Liu, X., Xu, J., Rosenthal, S., Zhang, L., McCubbin, R., Meshgin, N., Shang, L., Koyama, Y., Ma, H.-Y., Sharma, S., Heinz, S., Glass, C. K., Benner, C., Brenner, D. A., & Kisseleva, T. (2020). Identification of Lineage-Specific Transcription Factors That Prevent Activation of Hepatic Stellate Cells and Promote Fibrosis Resolution. Gastroenterology, 158(6), 1728-1744.e14. https://doi.org/10.1053/j.gastro.2020.01.027

Mak, K. M., & Png, C. Y. M. (2020). The Hepatic Central Vein: Structure, Fibrosis, and Role in Liver Biology. The Anatomical Record, 303(7), 1747–1767. https://doi.org/10.1002/ar.24273

Michalopoulos, G. K., & DeFrances, M. C. (1997). Liver Regeneration. Science, 276(5309), 60–66. https://doi.org/10.1126/science.276.5309.60

Monga, S. P. (2015). β-Catenin Signaling and Roles in Liver Homeostasis, Injury, and Tumorigenesis. Gastroenterology, 148(7), 1294–1310. https://doi.org/10.1053/j.gastro.2015.02.056

Mossanen, J., & Tacke, F. (2015). Acetaminophen-induced acute liver injury in mice. Laboratory Animals, 49(1_suppl), 30–36. https://doi.org/10.1177/0023677215570992

Nakamura, T., Nawa, K., & Ichihara, A. (1984). Partial purification and characterization of hepatocyte growth factor from serum of hepatectomized rats. Biochemical and Biophysical Research Communications, 122(3), 1450–1459. https://doi.org/10.1016/0006-291X(84)91253-1

Nakano, Y., Nakao, S., Sumiyoshi, H., Mikami, K., Tanno, Y., Sueoka, M., Kasahara, D., Kimura, H., Moro, T., Kamiya, A., Hozumi, K., & Inagaki, Y. (2017). Identification of a novel alpha-fetoprotein-expressing cell population induced by the Jagged1/Notch2 signal in murine fibrotic liver. Hepatology Communications, 1(3), 215–229. https://doi.org/10.1002/hep4.1026

Newman, A. M., Steen, C. B., Liu, C. L., Gentles, A. J., Chaudhuri, A. A., Scherer, F., Khodadoust, M. S., Esfahani, M. S., Luca, B. A., Steiner, D., Diehn, M., & Alizadeh, A. A. (2019). Determining cell type abundance and expression from bulk tissues with digital cytometry. Nature Biotechnology, 37(7), 773–782. https://doi.org/10.1038/s41587-019-0114-2

Otsu, N. (1979). A Threshold Selection Method from Gray-Level Histograms. IEEE Transactions on Systems, Man, and Cybernetics, 9(1), 62–66. https://doi.org/10.1109/TSMC.1979.4310076

Pellicoro, A., Ramachandran, P., Iredale, J. P., & Fallowfield, J. A. (2014). Liver fibrosis and repair: Immune regulation of wound healing in a solid organ. Nature Reviews Immunology, 14(3), 181– 194. https://doi.org/10.1038/nri3623

Planas-Paz, L., Orsini, V., Boulter, L., Calabrese, D., Pikiolek, M., Nigsch, F., Xie, Y., Roma, G., Donovan, A., Marti, P., Beckmann, N., Dill, M. T., Carbone, W., Bergling, S., Isken, A., Mueller, M., Kinzel, B., Yang, Y., Mao, X., … Tchorz, J. S. (2016). The RSPO–LGR4/5–ZNRF3/RNF43 module controls liver zonation and size. Nature Cell Biology, 18(5), 467–479. https://doi.org/10.1038/ncb3337

Poisson, J., Lemoinne, S., Boulanger, C., Durand, F., Moreau, R., Valla, D., & Rautou, P.-E. (2017). Liver sinusoidal endothelial cells: Physiology and role in liver diseases. Journal of Hepatology, 66(1), 212–227. https://doi.org/10.1016/j.jhep.2016.07.009

Puche, J. E., Saiman, Y., & Friedman, S. L. (2013). Hepatic Stellate Cells and Liver Fibrosis. Comprehensive Physiology, 3, 20.

Recknagel, R. O. (1967). Carbon Tetrachloride Hepatotoxicity. Pharmacological Reviews, 19(2), 145–208.

Remmerie, A., Martens, L., Thoné, T., Castoldi, A., Seurinck, R., Pavie, B., Roels, J., Vanneste, B., De Prijck, S., Vanhockerhout, M., Binte Abdul Latib, M., Devisscher, L., Hoorens, A., Bonnardel, J., Vandamme, N., Kremer, A., Borghgraef, P., Van Vlierberghe, H., Lippens, S., … Scott, C. L. (2020). Osteopontin Expression Identifies a Subset of Recruited Macrophages Distinct from Kupffer Cells in the Fatty Liver. Immunity, 53(3), 641-657.e14. https://doi.org/10.1016/j.immuni.2020.08.004

Ruoslahti, E., & Seppälä, M. (1979). α-Fetoprotein in Cancer and Fetal Development1. In G. Klein & S. Weinhouse (Eds.), Advances in Cancer Research (Vol. 29, pp. 275–346). Academic Press. https://doi.org/10.1016/S0065-230X(08)60849-0

Schiødt, F. V., Ostapowicz, G., Murray, N., Satyanarana, R., Zaman, A., Munoz, S., & Lee, W. M. (2006). Alpha-fetoprotein and prognosis in acute liver failure. Liver Transplantation, 12(12), 1776–1781. https://doi.org/10.1002/lt.20886

Schmidt, L. E., & Dalhoff, K. (2005). Alpha-fetoprotein is a predictor of outcome in acetaminophen-induced liver injury. Hepatology, 41(1), 26–31. https://doi.org/10.1002/hep.20511

Schwanhäusser, B., Busse, D., Li, N., Dittmar, G., Schuchhardt, J., Wolf, J., Chen, W., & Selbach, M. (2011). Global quantification of mammalian gene expression control. Nature, 473(7347), 337–342. https://doi.org/10.1038/nature10098

Shi, C., & Pamer, E. G. (2011). Monocyte recruitment during infection and inflammation. Nature Reviews Immunology, 11(11), 762–774. https://doi.org/10.1038/nri3070

Singh, S., Hynan, L. S., Rule, J. A., & Lee, W. M. (2019). Changes in alpha-foetoprotein and Gc-globulin in relation to outcomes in non-acetaminophen acute liver failure. Liver International: Official Journal of the International Association for the Study of the Liver, 39(12), 2368–2373. https://doi.org/10.1111/liv.14216

Tacke, F., & Zimmermann, H. W. (2014). Macrophage heterogeneity in liver injury and fibrosis. Journal of Hepatology, 60(5), 1090–1096. https://doi.org/10.1016/j.jhep.2013.12.025

Tanami, S., Ben-Moshe, S., Elkayam, A., Mayo, A., Bahar Halpern, K., & Itzkovitz, S. (2017). Dynamic zonation of liver polyploidy. Cell and Tissue Research, 368(2), 405–410. https://doi.org/10.1007/s00441-016-2427-5

Tipton, D. A., & Dabbous, M. K. (1998). Autocrine Transforming Growth Factor β Stimulation of Extracellular Matrix Production by Fibroblasts From Fibrotic Human Gingiva. Journal of Periodontology, 69(6), 609–619. https://doi.org/10.1902/jop.1998.69.6.609

Troeger, J. S., Mederacke, I., Gwak, G., Dapito, D. H., Mu, X., Hsu, C. C., Pradere, J., Friedman, R. A., & Schwabe, R. F. (2012). Deactivation of Hepatic Stellate Cells During Liver Fibrosis Resolution in Mice. Gastroenterology, 143(4), 1073-1083.e22. https://doi.org/10.1053/j.gastro.2012.06.036

Wang, G., Chen, S., Zhao, C., Li, X., Zhang, L., Zhao, W., Chang, C., & Xu, C. (2016). Gene expression profiles predict the possible regulatory role of OPN-mediated signaling pathways in rat liver regeneration. Gene, 576(2, Part 2), 782–790. https://doi.org/10.1016/j.gene.2015.11.008

Werb, Z., & Gordon, S. (1975). Elastase secretion by stimulated macrophages. Characterization and regulation. Journal of Experimental Medicine, 142(2), 361–377. https://doi.org/10.1084/jem.142.2.361

Xiong, X., Kuang, H., Ansari, S., Liu, T., Gong, J., Wang, S., Zhao, X.-Y., Ji, Y., Li, C., Guo, L., Zhou, L., Chen, Z., Leon-Mimila, P., Chung, M. T., Kurabayashi, K., Opp, J., Campos-Pérez, F., Villamil-Ramírez, H., Canizales-Quinteros, S., … Lin, J. D. (2019). Landscape of Intercellular Crosstalk in Healthy and NASH Liver Revealed by Single-Cell Secretome Gene Analysis. Molecular Cell, 75(3), 644-660.e5. https://doi.org/10.1016/j.molcel.2019.07.028

Yanger, K., Knigin, D., Zong, Y., Maggs, L., Gu, G., Akiyama, H., Pikarsky, E., & Stanger, B. Z. (2014). Adult Hepatocytes Are Generated by Self-Duplication Rather than Stem Cell Differentiation. Cell Stem Cell, 15(3), 340–349. https://doi.org/10.1016/j.stem.2014.06.003

Zhao, L., Jin, Y., Donahue, K., Tsui, M., Fish, M., Logan, C. Y., Wang, B., & Nusse, R. (2019). Tissue Repair in the Mouse Liver Following Acute Carbon Tetrachloride Depends on Injury-Induced Wnt/β-Catenin Signaling. Hepatology, 69(6), 2623–2635. https://doi.org/10.1002/hep.30563

Zigmond, E., Samia-Grinberg, S., Pasmanik-Chor, M., Brazowski, E., Shibolet, O., Halpern, Z., & Varol, C. (2014). Infiltrating Monocyte-Derived Macrophages and Resident Kupffer Cells Display Different Ontogeny and Functions in Acute Liver Injury. The Journal of Immunology, 193(1), 344– 353. https://doi.org/10.4049/jimmunol.1400574

